# Unrestrained Gα_i2_ Signaling Disrupts Normal Neutrophil Trafficking, Aging, and Clearance

**DOI:** 10.1101/2020.10.09.333112

**Authors:** Serena Li-Sue Yan, Il-Young Hwang, Olena Kamenyeva, Juraj Kabat, Ji Sung Kim, Chung Park, John H. Kehrl

## Abstract

Neutrophil trafficking, homeostatic and pathogen elicited, depends upon chemoattractant receptor triggered heterotrimeric G-protein Gα_i_βγ signaling, whose magnitude and kinetics are governed by RGS protein/Gα_i_ interactions. Yet how in totality RGS proteins shape neutrophil chemoattractant receptor activated responses remains unclear. Here, we show that C57Bl/6 neutrophils with genomic knock-in of a mutation that disables all RGS protein-Gα_i2_ interactions (G184S) cannot properly balance chemoattractant receptor signaling, nor appropriately respond to inflammatory insults. Mutant neutrophils accumulate in bone marrow, spleen, lung, and liver; despite neutropenia and an intrinsic inability to properly mobilize from bone marrow. *In vitro* they rapidly adhere to ICAM-1 coated plates, but poorly adhere to blood vessel endotheliums *in vivo*. Those few neutrophils that cross endotheliums migrate haphazardly. Following Concanavalin-A administration fragmented G184S neutrophils accumulate in liver sinusoids leading to thrombo-inflammation and perivasculitis. Thus, neutrophil Gα_i2_/RGS protein interactions limit and facilitate Gα_i2_ signaling allowing normal neutrophil trafficking, aging, and clearance.

## Introduction

Signaling via CXCR2 and CXCR4 plays several essential roles in neutrophil trafficking. CXCR2 and CXCR4 counter regulate the release of mature neutrophils from the bone marrow (BM) into the circulation (Eash et al., 2010). Either excessive CXCR4 signaling or a lack of CXCR2 signaling causes myelokathexis (Eash et al., 2010; Liu et al., 2015). Myelokathexis is the inappropriate retention (kathexis) of neutrophils (myelo) in the BM. CXCR2 signaling also triggers diurnal changes in the transcriptional and migratory properties of circulating neutrophils, a process termed neutrophil aging (Adrover et al., 2020; Adrover et al., 2019). These diurnal changes are opposed by CXCR4 signaling. Neutrophils released from the BM express high levels of CD62L that progressively decline during the day, while increasing amounts of surface CXCR4 promotes neutrophil egress from the blood into the tissues, which leads to their clearance by tissue macrophages (Adrover et al., 2019; Adrover et al., 2016). Deletion of CXCR2 from neutrophils prevents phenotypic aging, whereas deletion of CXCR4 promotes unrestrained aging (Adrover et al., 2019). Neutrophil aging impairs CD62L mediated endothelial rolling, thereby reducing neutrophil accumulation at sites of inflammation (Adrover et al., 2019). Yet aged neutrophils efficiently cross the endothelium to enter tissues. The removal of aging neutrophils from the circulation facilitates their clearance, helping to protect blood vessels from potential neutrophil mediated insults (Adrover et al., 2019; Hidalgo et al., 2019).

Both CXCR2 and CXCR4 as well as other chemoattractant receptors that help control leukocyte and neutrophil recruitment and trafficking use Gαi to link to downstream effector molecules (Eash et al., 2010; Furze and Rankin, 2008; Kehrl, 2006). Ligand engagement of chemoattractant receptors trigger a conformational change that facilitates receptor/heterotrimeric G-protein coupling, Gαi subunit GDP-GTP exchange, functional Gαi dissociation from Gβγ subunits, downstream effector activation leading to integrin activation, and directed migration (Cho and Kehrl, 2009; Kamp et al., 2016). Since Gαi subunits possess an intrinsic GTPase activity, GTP hydrolysis facilitates re-assembly of the heterotrimeric G-protein causing signaling to cease (Cho and Kehrl, 2009; Kehrl, 2016). By dramatically accelerating the intrinsic GTPase activity of Gαi subunits, RGS proteins reduce the duration that Gαi subunits remains GTP bound, thereby decreasing effector activation by reducing available Gαi-GTP and free Gβγ (Kehrl, 2016).

Despite the importance of Gα_i_-coupled receptors in neutrophil trafficking and aging, relatively little is known about the overall importance of RGS proteins in chemoattractant receptor signaling. Murine neutrophils prominently express *Rgs2, Rgs18*, and *Rgs19;* lesser amounts of *Rgs3* and *Rgs14;* and detectable levels of mRNA transcripts for several other RGS proteins (http://www.immgen.org/databrowser/index.html). They also highly express *Gnai2*, with a lower amount of *Gnai3* (approximately 1/5 the amount as assessed by RNA sequencing) and little or no *Gnai1* or *Gnao*. Loss of *Rgs2* in mice increases neutrophil recruitment to inflamed lungs (George et al., 2017; George et al., 2018). Despite its low expression level, loss of *Rgs5* in mice also leads to a more robust recruitment of neutrophils to inflamed lungs (Chan et al., 2018). In addition, purified neutrophils from these mice had exaggerated responses to CXCR2 and CXCR4 ligands (Chan et al., 2018). Since the loss an of individual RGS proteins often causes a relatively mild phenotype, perhaps a consequence of their redundant expression profiles, we have made use of a genetically modified mouse that has a mutation, which replaces glycine 184 in the Gα_i2_ protein with a serine (Huang et al., 2006; Lan et al., 1998). This change blocks the binding of RGS proteins to Gα_i2_, thereby rendering RGS proteins unable to act as GTPase activating proteins (GAPs). We will refer to mice or neutrophils bearing the Gα_i2_ G184S mutation on both alleles as G184S mice or neutrophils, respectively. Previously, we have shown that the G184S mice accumulate BM neutrophils, which mobilize poorly to an inflamed peritoneum or in response to sterile ear inflammation. Also, these mice did not control a normally nonlethal *Staphylococcus aureus* infection (Cho et al., 2012).

In this study we have further investigated the trafficking patterns and mobilization of G184S neutrophils to inflammatory stimuli predominately using mice reconstituted with either WT or G184S BM, or with a 1:1 mixture. We relied on intravital microscopy to image the behavior of WT and mutant neutrophils in the BM, lung, lymph node, and liver. Our studies indicate the G184S Gα_i2_ mutation causes an initial misbalance in the BM between the CXCR4 mediated retention signal and the CXCR2 mediated mobilization signal. Those neutrophils that escape the BM rapidly leave the circulation to accumulate in the marginal pools located in the lung, liver, and spleen. Neutrophil recruitment to inflammatory sites is severely impaired secondary to both a mobilization defect and to impaired transendothelial migration (TEM). Furthermore, G184S BM reconstituted mice poorly tolerate ConA induced inflammation as G184S neutrophils fragment and aggregate in liver sinusoids. The significant of these results are discussed.

## Results

### Neutrophil Overload in G184S BM Reconstituted Mice Despite Neutropenia

We determined the numbers of neutrophils in the blood, BM, and spleen of mice reconstituted with WT or G184S BM. Neutrophils lack the lineage markers B220, CD3, CD4, CD8, CD11c, NK1.1, c-kit, and TCRγδ, but express CD11b and Ly6G. We assessed neutrophil maturity by the level of CD11b expression on Ly6G^+^ cells (Casanova-Acebes et al., 2013; Hidalgo et al., 2019). Representative flow cytometry patterns and the numbers of leukocytes and neutrophils at the different sites are shown. The G184S BM reconstituted mice have a reduced number of blood neutrophils with expanded populations in the spleen, lung, and the BM (Figure 1A). The spleen had a 10-fold, the lung an 8-fold, and the BM a 2-fold expansion of neutrophils compared to WT while the blood had a 3.5-fold reduction. The G184S BM reconstituted mice also had a relative increase in mature neutrophils in the BM (CD11b^high^ and Ly6G^+^) compared to WT BM reconstituted mice, while they had an increase of immature cells (CD11b^low^Ly6G^+^) in the spleen (Figure 1B).

**Figure 1.**
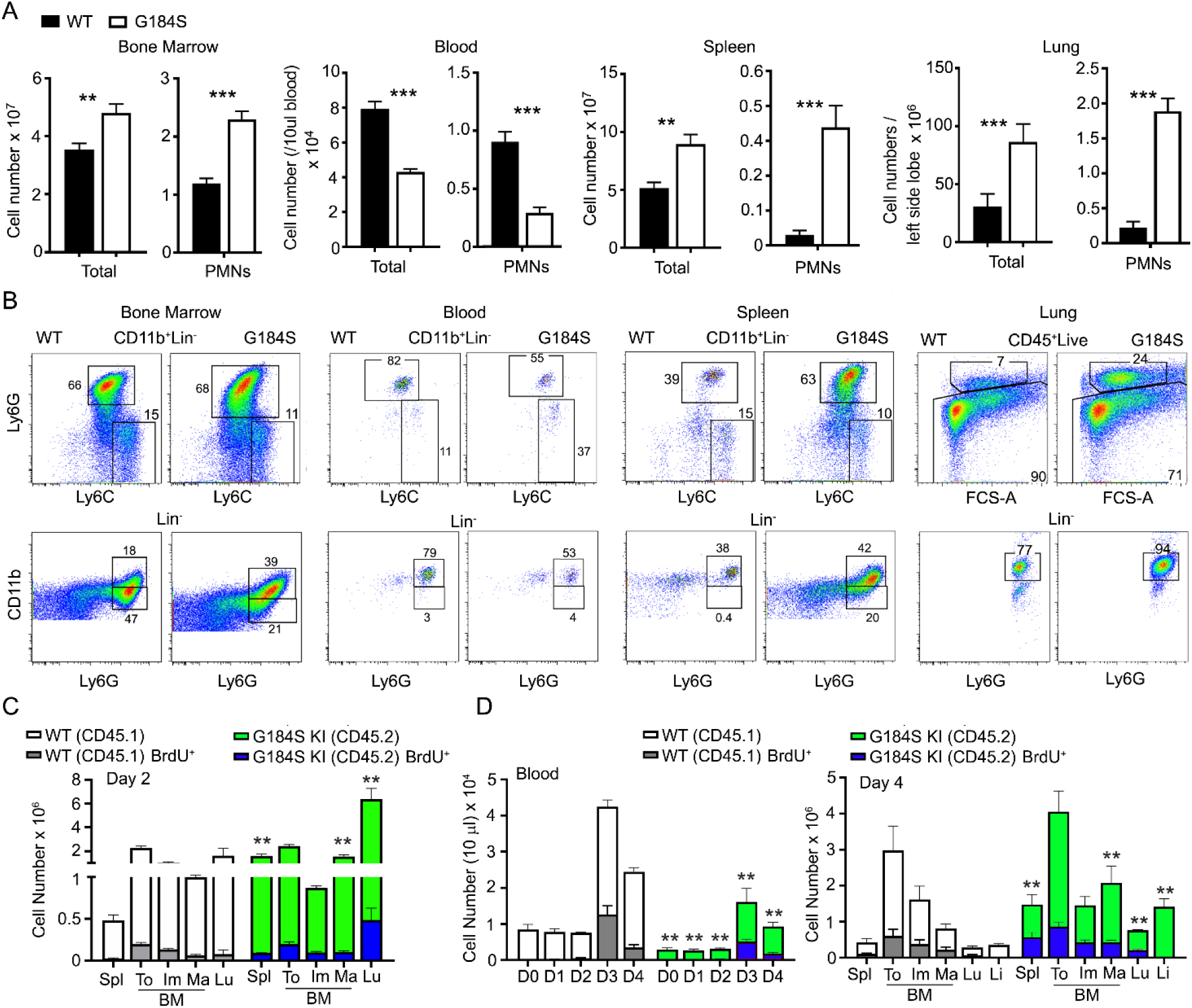
Analysis of Neutrophil distribution in WT and G184S BM Reconstituted Mice. (**A**) A comparison of total leukocyte and neutrophil (PMN) blood cell counts between WT and G184S BM (BM) reconstituted mice in BM, blood, spleen, and lung. Neutrophils are identified as CD11b^+^/Ly6G^+^ cells by flow cytometry. Results are from 3 separate experiments done in triplicates from using WT (n = 3) and G184S (n = 3) BM reconstituted mice. PMNs, neutrophils. Statistics: data are means ± SEMs and analyzed using unpaired Student’s *t*-test comparing G184S with WT. (**B**) Representative flow cytometry patterns and numbers of leukocytes and neutrophils at BM, blood, spleen, and lung are shown. (**C**) Cell count comparison of WT vs G184S neutrophil production rate and distribution 2 days after BrdU injection in 1:1 (WT:G184S) mixed chimeras. Spl, spleen; To, total neutrophils; Im, immature; Ma, mature; Lu, lung. (**D**) Cell count comparison of WT vs G184S neutrophil production rate and distribution 2 days after BrdU injection and CD62L antibody treatment in mixed chimeras. Graph on the left shows the neutrophil count in blood from days 0 to 4 (D0 – D4), with BrdU and CD62L antibody administered on D2. Graph on the right shows the WT vs G184S neutrophil production rate and distribution on D4. 1(**A, C, D**) Statistics: data are means ± SEMs, and analyzed using unpaired Student’s *t*-test comparing G184S with WT. (**C, D**) Results are from 3 separate experiments done in duplicates from 3 mixed chimeras for each study. *p < 0.05, **p < 0.005 and ***p < 0.0005

The neutropenia in the setting of an expanded BM population of mature neutrophils suggests a BM egress defect. Yet a BM egress defect can’t explain the peripheral neutrophil expansion noted in these mice suggesting a more complex phenotype. To assess whether the concomitant presence of WT neutrophils would normalize the G184S neutrophil distribution we made mixed BM chimeric mice. C57Bl/6 CD45.2 mice were reconstituted with 50% CD45.2 WT BM and 50% CD45.1 G184S BM. After 6-8 weeks we assessed the number of neutrophils in various locations using CD45.1 and CD45.2 monoclonal antibodies to distinguish the two genotypes. We performed two sets of experiments in the first we injected the mixed chimeric mice with BrdU two day prior to sacrificing the mice for analysis (Figure 1C). In the second we again injected BrdU, but on day 2 we treated the mice with a CD62L antibody to reduce neutrophil transendothelial migration and margination; and sacrificed the mice on day 4 (Figure 1D). In sum, the mixed chimera BM contained a nearly equal ratio of WT and G184S neutrophils, although the G184S cells predominated among the mature BM neutrophils. The WT neutrophils exceeded the G184S cells in the blood, 70% versus 30%, while the lung, liver, and spleen each contained a 3 to 4-fold excess of G184S versus WT neutrophils. The CD62L treatment raised the blood neutrophil counts causing a 5 to 6-fold increase compared to basal for both the WT and G184S cells. The BrdU labeling suggested that the neutrophil production rates for the two genotypes did not differ appreciably (Figure 1D). The coexistence of the WT and G184S neutrophils in the mixed chimeric mice partially corrected the BM expansion, failed to correct the neutropenia, and partially corrected the peripheral neutrophil overload noted in the G184S straight chimeric mice. These results suggest that the neutropenia in the G184S mice results not only from an egress defect, but also from a short intravascular half-life. The loss of RGS protein/Gα_i2_ interactions results in expanded pools of neutrophils in spleen, lungs, and the liver.

### Localization of Bone Marrow and Peripheral G184S Neutrophils

To address the localization of the G184S neutrophils in the BM, spleen, lung, and liver we used a combination of immunohistochemistry and intravital microscopy. For these experiments, we relied on G184S and WT BM reconstituted mice, or occasionally on non-reconstituted WT and G184S mice. Two-photon microscopy of the skull BM confirmed the expanded neutrophil BM niche in the G184S mice. Mice with the LysM-GFP transgene crossed onto a G184S background were used for the BM imaging. We visualized the BM vasculature by infusing Evans blue dye into the bloodstream. A stitched set of 2-photon images spanning the BM niches are shown. Higher magnification images show the high neutrophil density in the neutrophil niches of the G184S BM (Figure S1A). Next, we used confocal microscopy to examine fixed spleens from mice reconstituted with WT or G184S BM and immunostained with CD169 to delineate the splenic marginal zone and the separation between the white pulp and red pulp; and Ly-6G to identify neutrophils. Stitched confocal images show the larger spleen size and the markedly expanded neutrophil population in the splenic red pulp of G184S BM reconstituted mice. (Figure S1B). Higher magnification inserts demonstrate the infiltration of G184S neutrophils into the white pulp. Normally, in young mice neutrophils do not enter the white pulp or lymph node follicles (Kamenyeva et al., 2015; Tomay et al., 2018). As mice age, rare neutrophils can be found in the white pulp, and infection or LPS challenge can also cause neutrophils to enter the white pulp or lymph node follicles (Bronte and Pittet, 2013; Kamenyeva et al., 2015). Live confocal microscopy of lung sections immunostained with CD31 and Gr-1 antibodies confirmed the marked neutrophil expansion in the lungs of the G184S BM reconstituted mice (Figure S1C). CD31 immunostaining delineated the pulmonary endothelial cells and outlined the pulmonary vasculature. Higher magnification inserts show that the G184S neutrophils largely resided within the pulmonary vasculature. Finally, we used intravital confocal microscopy to image the liver outlining the liver sinusoids by injection of labeled CD31 and the neutrophils by injecting labeled Gr-1 antibody. At baseline we found relatively few neutrophils in the superficial sinusoids of the WT BM reconstituted mice livers while we consistently observed neutrophils in the liver sinusoids of the G184S reconstituted mice (Figure S1D). These data confirm the expanded BM niche and demonstrates that the G184S neutrophils accumulate in the splenic red pulp, lung vasculature, and liver sinusoids. These results suggest that the loss of Gα_i2_/RGS protein interactions in neutrophils dramatically expands the marginated pool of neutrophils.

### Neither CXCL1 nor AMD3100 Efficiently Mobilizes G184S Neutrophils to the Blood, but both Signals Do, as Does KRH3955

We had previously found that peritoneal inflammation did not efficiently recruit G184S neutrophils into the bloodstream (Cho et al., 2012). To better understand the mobilization defect we tested whether raising CXCL1 blood levels (Girbl et al., 2018), would mobilize the G184S BM neutrophils (Figure 2A). Injection of WT mice increased blood neutrophils 8-fold, while blood G184S neutrophils increased, they did not reach the basal level found in WT mice. Next, we tried the CXCR4 antagonist AMD3100, which increases blood neutrophil numbers perhaps by reversing the BM CXCL12 gradient (Pillay et al., 2020; Redpath et al., 2017). Again, while AMD3100 treatment increased the blood WT neutrophils, the blood neutrophil numbers in G184S mice did not reach the basal level found in WT mice (Figure 2A). We also assessed the impact of CXCL1 and AMD3100 administration on the bone marrow niche using intravital imaging. Initial comparison one hour after PBS injection revealed the increased density of LysM-GFP expressing cells in the G184S BM as noted previously, but also an increased basal motility of the cells within the BM niche (Figure 2B, Video S1). In both sets of mice, we observed relatively few LysM-GFP positive cells in the BM vasculature. The administration of CXCL1 or AMD3100 increase the number of neutrophils in the vasculature, more evident with CXCL1 than AMD3100, and in the WT versus the G184S mice. Both AMD3100 and KC increased the motility of WT BM neutrophils without obviously affecting the enhanced motility of the G184S neutrophils (Figure 2B, Video S2). Together these results reveal an increase in the basal motility of G184S neutrophils along with an underlying defect in their ability to be mobilized to the blood by raising CXCR2 signaling or by reversing the CXCL12 gradient.

**Figure 2.**
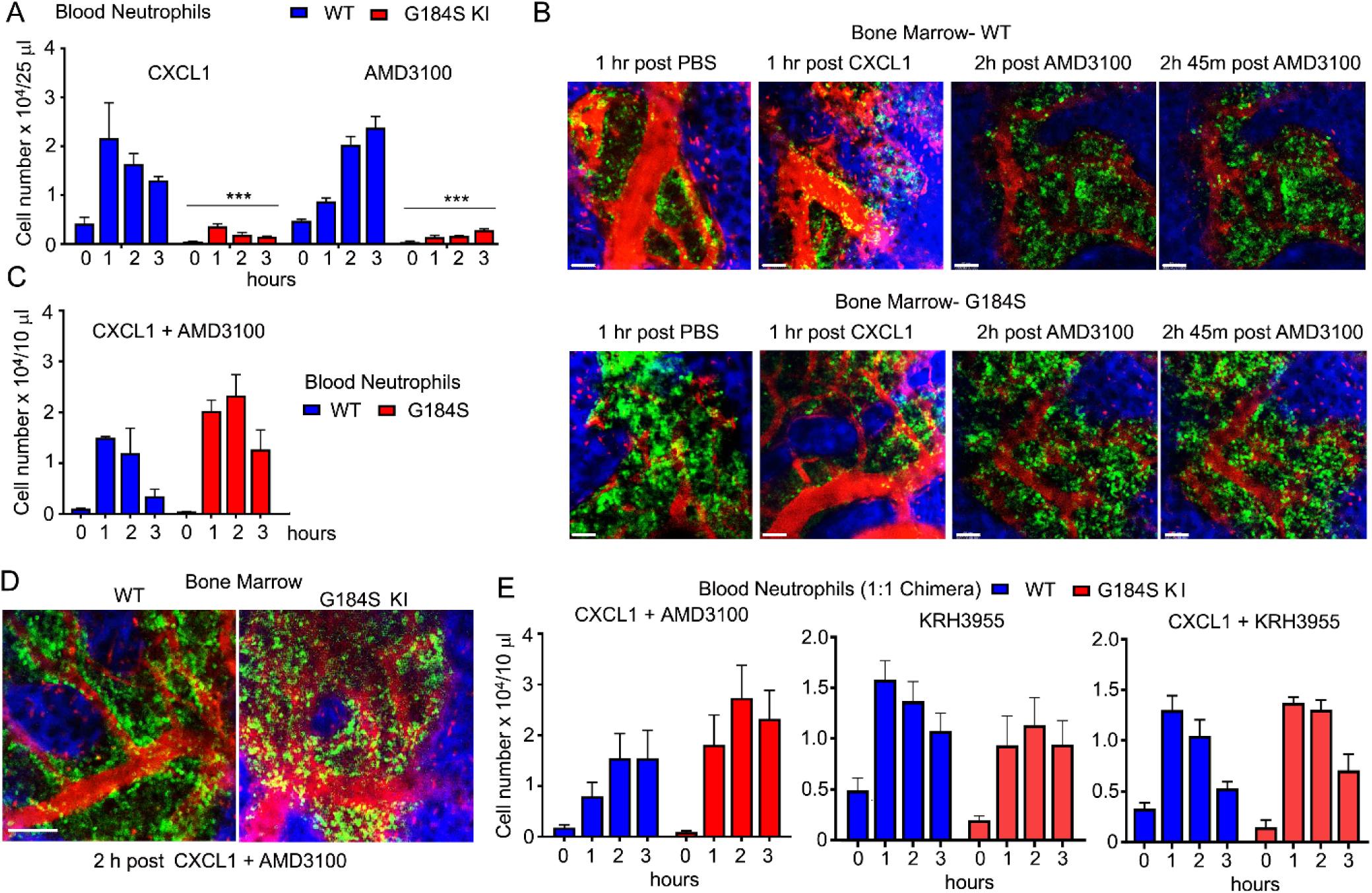
Effects of CXCL1, AMD3100, and KRH3955 on WT vs G184S Neutrophil Mobilization. (**A**) Circulating blood neutrophil counts in WT vs G184S BM reconstituted mice after injections of CXCL1 (i.v.) or AMD3100 (i.p.). Blood neutrophil counts were monitored from 0 (basal) – 3 hrs after CXCL1 or AMD3100 injections. (**B**) Representative 2P-IVM images of skull BM in WT vs G184S mice with LysM-GFP neutrophils (green) and Evans Blue labeled blood vessels (red). Comparison of WT (top panels) to G184S (bottom panels) neutrophil BM localization and intravascular mobilization after either PBS (control), CXCL1, or AMD3100 injections. Scale bar = 50 µm. (**C**) Levels of circulating blood neutrophils in WT vs G184S BM reconstituted mice after injections of both CXCL1 (i.v.) and AMD3100 (i.p.) 20 min apart. Blood neutrophil counts were monitored from 0 (basal) – 3 hrs after CXCL1 + AMD3100 injections. (**D**) Representative 2P-IVM images of skull BM in WT vs G184S mice with LysM-GFP neutrophils (green) and Evans Blue outlined blood vessels (red). Comparison of WT (left) to G184S (right) BM neutrophil localization and intravascular mobilization 2 hrs after CXCL1 and AMD3100 injections. Scale bar = 100 µm. (**E**) Circulating blood neutrophil counts in WT vs G184S 1:1 BM reconstituted mice after injection of either CXCL1 with AMD3100 (left), KRH3955 alone (middle), or CXCL1 with KRH3955 (right). Blood neutrophil counts were monitored from 0 (basal) – 3 hrs after each treatment. (**A, C**) Results are from 3 separate experiments done in duplicates from WT (n = 3) and G184S (n = 3) BM reconstituted mice for each study. Statistics: data are means ± SEMs and analyzed using unpaired Student’s *t*-test comparing G184S with WT. (**B, D**) Images are representative of WT (n = 3) and G184S (n = 3) mice for each study. (**E**) Results are from 3 separate experiments done in duplicates from 3 1:1 (WT:G184S) mixed chimeras. ***p < 0.0005

To provide a stronger mobilization signal, we co-administered AMD3100 and CXCL1. When given simultaneously both the WT and G184S mice rapidly died, however when staggered, AMD3100 followed 20 minutes later by CXCL1, the mice suffered no apparent ill effects. The staggered agents raised the blood level of WT neutrophils approximately 14-fold, nearly two-fold higher than either alone. The G184S neutrophils increased 40-fold above the basal exceeding the number of neutrophils mobilized in WT mice (Figure 2C). Intravital microscopy of BM revealed an increase in intravascular neutrophils in both the WT and G184S BMs (Figure 2D). We repeated the co-administration experiment using 1:1 mixed BM chimeric mice. Treatment of with AMD3100 and CXCL1 led to a 10-fold increase and 40-fold increase in the WT and G184S blood neutrophils, respectively (Figure 2E). We also tested a second CXCR4 antagonist, which does not reverse the CXCL12 BM gradient, but acts as a true CXCR4 antagonist (Redpath et al., 2017). Surprisingly, KRH3955 efficiently mobilized both WT and G184S neutrophils without having to co-administer CXCL1 (Figure 2E). These data indicate the loss of Gα_i2_/RGS protein interactions causes a misbalance between the CXCR4-mediated retention signal and the CXCR2-mediated recruitment in the BM. They also show that normal neutrophil BM egress depends upon RGS proteins limiting Gα_i2_ signaling.

### Abnormal G184S Neutrophil Aging

Once neutrophils are released from the BM, neutrophil aging begins. Over time CXCR4 levels rise and CXCR2 and CD62L levels decline. Aged and “fresh” neutrophils predominate in the blood at Zeitgeber time 5 (ZT5, 5 hours after lights are on) and ZT13, respectively (Casanova-Acebes et al., 2013). These changes in expression affect neutrophil BM release, recruitment to inflammatory sites, and peripheral clearance. To determine if the G184S neutrophils aged appropriated, we first assessed the above receptors on neutrophils collected from different sites using the 1:1 chimeric mice. The G184S BM neutrophils had higher CXCR4 membrane expression, more evident on the mature BM G184S neutrophils. The G184S BM neutrophils also had lower CXCR2 levels, which was accompanied by increased intracellular CXCR2 (Table S1). The G184S lung, liver, and splenic neutrophils all had significantly reduced levels of CXCR2 and CD62L; but their CXCR4 expression varied, slightly depressed on G184S lung neutrophils and slightly elevated on liver and splenic neutrophils (Table S2). Overall, the residential G184S neutrophils had slightly higher membrane CXCR4 levels, and lower amounts of CXCR2 and CD62L finding suggestive of a more aged phenotype.

Next, we checked for appropriate age-related changes by collecting blood at ZT5 and ZT13 for analysis. The G184S neutropenia and monocytopenia were present at both ZT5 and ZT13 (Figure 3A). At each time point the G184S blood neutrophils had higher CXCR4 and lower CXCR2 extracellular expression levels compared to WT neutrophils (Figure 3B). Surprisingly, we noted that in the mixed chimera mice, the WT neutrophils had a slightly higher extracellular CXCR2 expression level at ZT5 than at ZT13, opposite of the reported expression variation (Adrover et al., 2019). Total WT blood neutrophil CXCR2 expression (intracellular plus extracellular) did not differ between the time points, while the G184S neutrophils had a higher total CXCR2 expression level at ZT13 compared to ZT5. Compared to the WT cells ZT5 G184S neutrophils had elevated CD62L expression, and the ZT13 G184S neutrophils had even higher levels (Figure 3C). The shape changes that normally accompanies neutrophil aging did not occur in the G184S neutrophils (Figure 3D) and the ZT5 G184S neutrophils had reduced amounts of cortical actin and low pERM, consistent with microvilli loss (Adrover et al., 2019) (Figure 3E). Thus, while G184S blood neutrophil numbers exhibited a normal diurnal variation, their expression pattern of homing receptors, cell shape changes, and cortical actin levels did not conform to the aging patterns observed with the WT neutrophils.

**Figure 3.**
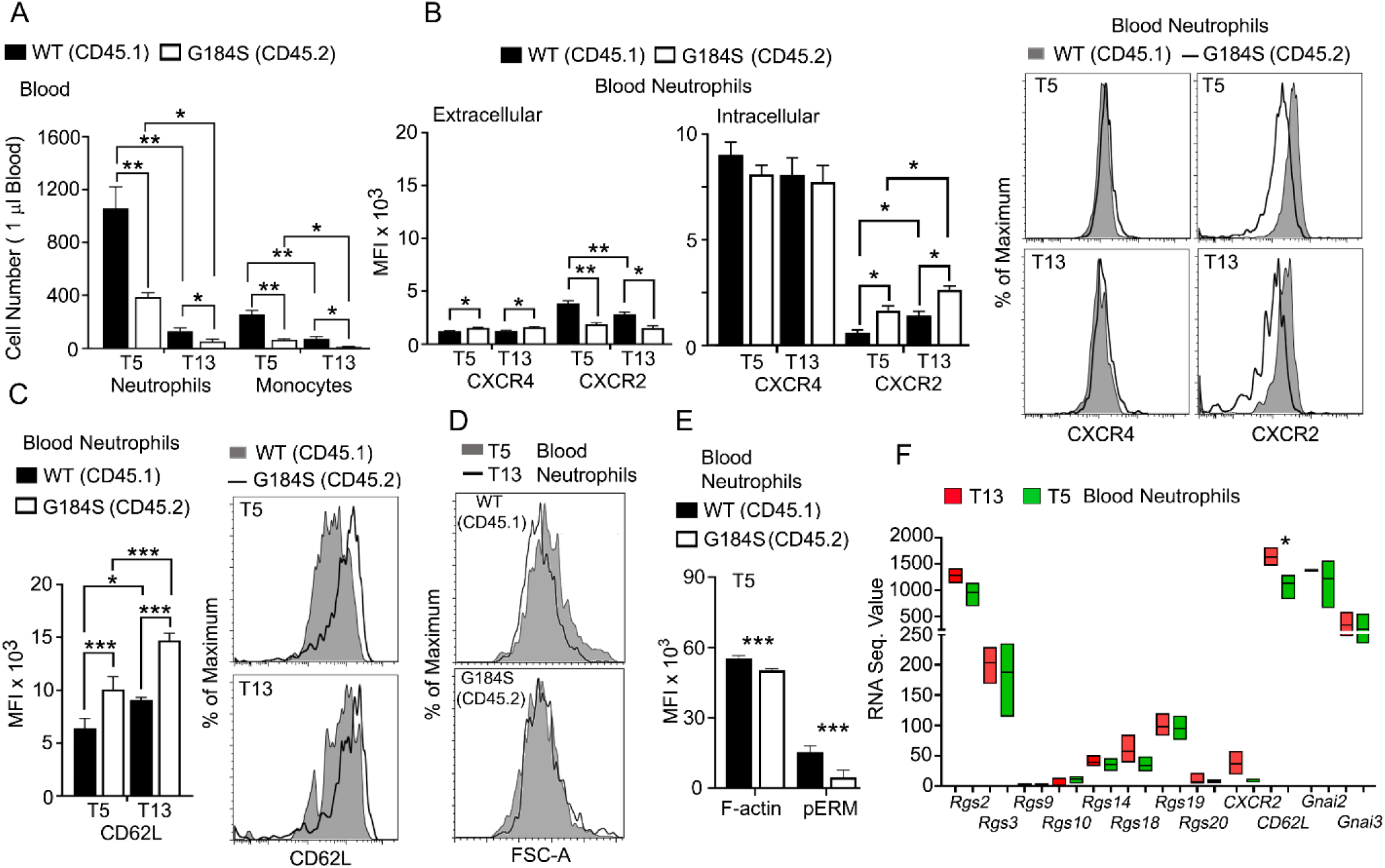
G184S Neutrophils Undergo Abnormal Aging with Altered CXCR4 and CXCR2 Expression Levels. (**A**) Circulating WT vs G184S blood neutrophil and monocyte counts at daytime (T5) and nighttime (T13). (**B**) Left graph, flow cytometry analysis of surface CXCR4 and CXCR2 expressions of WT vs G184S blood neutrophils at T5 (aged) and T13 (fresh). Middle graph, analysis of intracellular CXCR4 and CXCR2 levels. For intracellular CXCR4 and CXCR2 expression, cells were fixed, permeabilized, and stained for each chemokine receptor. Since surface chemokine receptors were not blocked prior to intracellular staining, intracellular chemokine receptor levels represent their total levels of the cell. Data are expressed as mean fluorescence intensities (MFI). Right, representative plots showing surface CXCR4 and CXCR2 expression on WT vs G184S blood neutrophils at T5 and T13. (**C**) MFI data (left graph) obtained from flow cytometry analysis of surface CD62L levels of WT vs G184S blood neutrophils at T5 and T13. Representative plots showing surface CD62L expression on WT vs G184 blood neutrophils AT T5 and T13 (right). (**A, B, C**) Results are from 3 separate experiments done in duplicates from 3 1:1 (WT:G184S) mixed chimeras for each study. Statistics: data are means ± SEMs analyzed by 1-way ANOVA. (**D**) Representative flow cytometry result comparing WT and G184S blood neutrophil forward light scatter. Blood neutrophils collected from 1:1 chimeric mice at ZT5 or ZT13 analyzed for forward light scatter. (**E**) Flow cytometry analysis of cortical actin and pEzrin levels of WT vs G184S blood neutrophils at T5. Statistics: data are means ± SEMs, and analyzed using unpaired Student’s *t*-test comparing G184S with WT. Results are from 3 1:1 (WT:G184S) BM reconstituted mice. (**F**) Relative mRNA expression data obtained from RNA seq experiments. RNA sequence value (arbitrary unit) comparing *Rgs* proteins, *Cxcr2, Cd62l, Gnai2*, and *Gnai3* mRNA expression levels of WT blood neutrophils at both T5 and T13. Statistics: data are means ± SEMs, then analyzed using unpaired Student’s *t*-test comparing T5 with T13. *p < 0.05, **p < 0.005 and ***p < 0.0005.

To determine whether changes in RGS protein expression accompany normal neutrophil aging, we checked previously published RNA sequence data (Adrover et al., 2020). We found that “aged” neutrophils tended to have lower levels of RGS protein expression, however the differences did not reach statistical significance (Figure 3F). *Gnai2* and *Gnai3* mRNA expression levels remained stable, although ZT13 cells had higher *Gnaq* mRNA levels (223 vs. 130, p < 0.02). Together these data indicate that changes in Gα_i_ protein expression is unlikely to impact neutrophil aging, while RGS2 warrants additional study.

### Acute Ligand Exposure Appropriately Downregulates CXCR4 and CXCR2 Expression

Besides age related changes ligand exposure can impact chemoattractant expression levels. Using the 1:1 mixed chimeric mice, we assessed the changes in CXCR4 and CXCR2 expression following *in vivo* administration of KRK3955, and CXCR2 expression following exposure to different ligands *in vitro*. Following KRH3955 administration membrane CXCR2 declined on both the WT and G184S PMNs, although the G184S neutrophil expression declined more gradually (Figure S2A). Since the WT blood neutrophils had a higher initial expression level, by three hours post KRH3955 no significant difference remained. The blood G184S neutrophil higher membrane CXCR4 levels declined gradually in parallel with the WT neutrophils after KRH3955 injection (Figure S2B). Overall, the % decline in CXCR4 and CXCR2 were not significantly different (Figure S2C). *In vitro* exposure to CXCL1, CXCL2, or to the chemoattractant fMLP led to similar declines in the WT and G184S neutrophil CXCR2 expression (Figure S2D & E). Overall, the loss of Gα_i2_/RGS protein interactions affected both CXCR2 and CXCR4 expression and signaling (see below), which in-turn likely impacted neutrophil aging.

### G184S Neutrophils Have Defects in *in vitro* TEM and Chemotaxis but Enhanced Basal Adhesion

Next, we examined whether the G184S neutrophil have a TEM defect using a transwell filter coated with the murine endothelial cells line (SVEC4-10). Freshly isolated BM–derived neutrophils were loaded on the apical side of the endothelial monolayers, and CXCL1 added to the bottom chamber. While the WT neutrophils readily crossed the endothelial cell barrier the G184S neutrophil largely failed (Figure 4A). We visualized the difficulty G184S PMNs had in crossing the endothelial barrier during the migration assay by differentially labeling the two sources of neutrophils and imaging across the cell barrier. The G184S neutrophils poorly penetrated the barrier appearing to be stuck on the endothelial cells (Figure 4A). Using standard transwell migration assays with non-coated filters we checked the migratory capacity of the WT and G184S neutrophils. Significant numbers of immature and mature G184S BM neutrophils migrated across the transwell barrier without added chemoattractant (Figure 4B). Despite this high basal migration, CXCL12 elicited comparable levels of specific neutrophil migration, while CXCL2 did not. The G184S BM neutrophil also migrated to sphingosine-1 phosphate, even exceeding that of WT cells (Figure 4B). Immature BM neutrophils migrated comparably to fMLP and C5a, while mature G184S BM neutrophils showed significant defects (Figure 4C). The specific migratory responses of G184S splenic neutrophils to CXCL2 and CXL12 were poor (Figure 4D).

**Figure 4.**
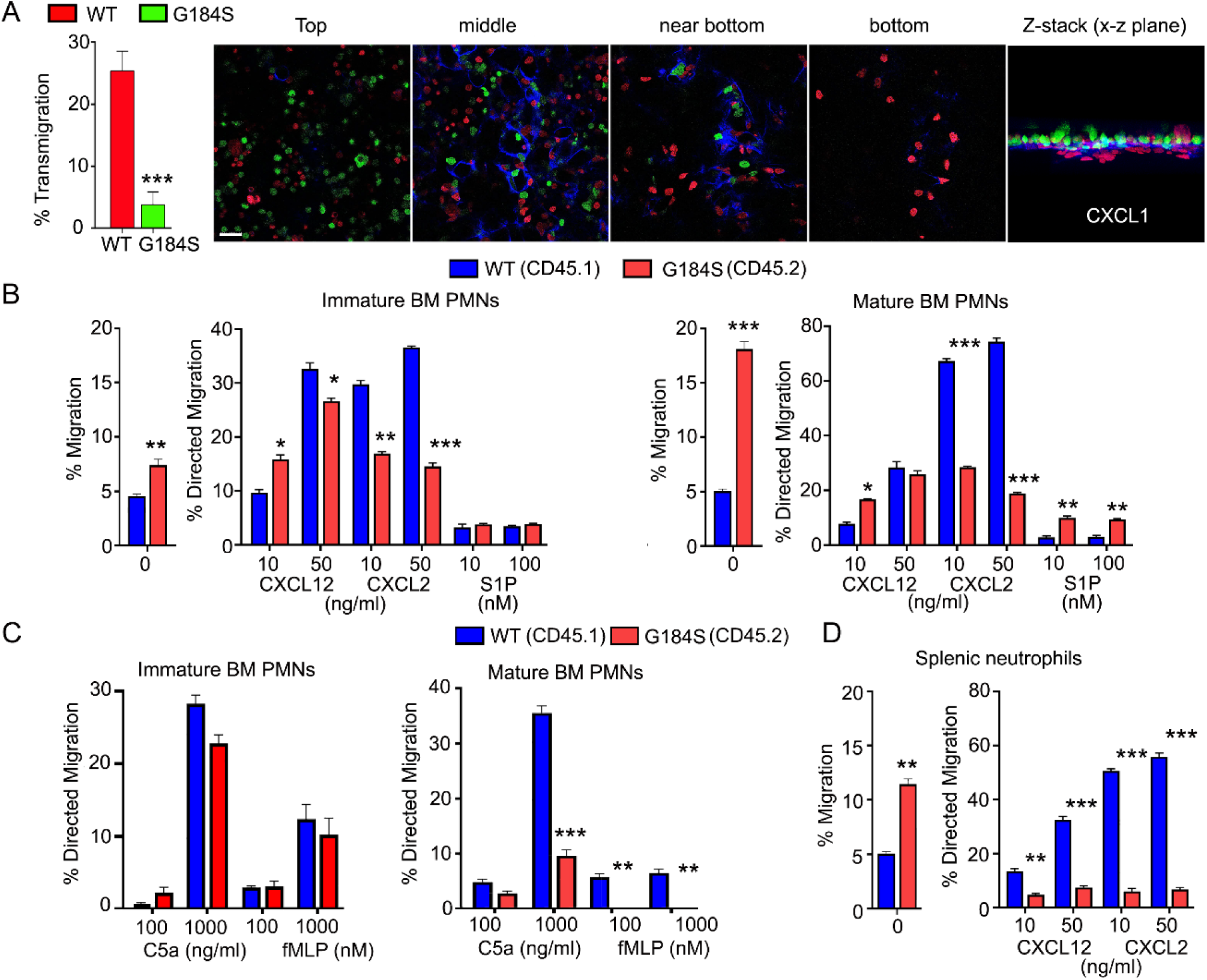
*In Vitro* TEM and Migratory Capacity of WT vs G184S Neutrophils. (**A**) Transmigration frequency (%) of WT vs G184S BM neutrophils to CXCL1 stimuli *in vitro* across SVEC4-10 coated transwell filters (left graph). Results are from 3 separate experiments done in duplicates. Representative confocal images (right panels) show the extent of differentially labeled WT (red) vs G184S (green) BM neutrophils crossing the endothelial barrier (blue) during the migration assay (top, middle near bottom, bottom). The Z-stack image viewed from the x-z plane (far right) shows majority of the WT neutrophils have crossed or in process of crossing the endothelial barrier, while the G184S neutrophils remain accumulated on the apical side of the barrier. Scale bar = 30 µm. Results are from BM neutrophils isolated from WT (n = 3) and G184S (n = 3) BM reconstituted mice. (**B**) *In vitro* chemotaxis assays assessing the migratory capacities of WT vs G184S immature (2 left graphs) and mature (2 right graphs) BM neutrophils with increasing concentrations of CXCL12, CXCL2 and S1P. Shown are the percentages that specifically migrated, with controls as ‘0’ treatment. (**C**) Chemotaxis assays assessing migratory capacities of WT vs G184S immature (left) and mature (right) BM neutrophils with increasing concentrations of C5a and fMLP *in vitro*. (**D**) Chemotaxis assays with increasing concentrations of CXCL12 and CXCL2 to assess migratory capacities of WT vs G184S splenic neutrophils *in vitro*. (**B, C, D**) Results are from 3 1:1 (WT:G184S) BM reconstituted mice with n ≥ 3 experiments done in duplicates for each study. Statistics: data are means ± SEMs, then analyzed using unpaired Student’s *t*-test comparing G184S with WT. *p < 0.05, **p < 0.005 and ***p < 0.0005.

As CXCR2 triggered migration showed the most severe impairment, we tested proximal CXCR2 signaling by measuring the intracellular calcium responses following sequential exposure to CXCL1 and CXCL2, which have been shown to mediate neutrophil TEM in vivo (Girbl et al., 2018). At baseline the G184S neutrophils had elevated intracellular calcium levels (Figure 5A). Relative to WT neutrophils, both CXCL1 and CXCL2 elicited substandard increases in intracellular calcium in the G184S neutrophils (Figure 5B). The CXCL1 response being more severely impaired. We also assessed CXCR2 triggered adhesion. Even in the absence of added chemokine, 11-fold more G184S neutrophils attached and spread on the ICAM-1 coated plates than did WT cells (Figure 5C). Pre-exposure to CXCL2 and MgCl_2_ prior to plating enhanced 4-fold the number of spread WT neutrophils. Due to the high background adhesion a similar pre-exposure minimally increased the spreading of G184S neutrophil. A comparison of footprint sizes of the pre-exposed neutrophils revealed a dramatic increase among the G184S neutrophils compared to the simultaneously assessed WT neutrophils (Figure 5C). Thus, basally the G184S neutrophils possess elevated intracellular calcium levels, increased motility, and enhanced adhesion to ICAM-1 coated plates. They also exhibit reduced specific migratory and calcium responses to CXCL1/2. Surprisingly, while C5a and fMLP elicit suboptimal migration, they triggered enhanced intracellular calcium responses (Figure 5D). This argues that the proximal signaling pathway leading to calcium mobilization following Gα_i_ activation is intact. Thus, the loss of Gα_i2_/RGS protein interactions enhances basal Gα_i2_ signaling while variably affecting ligand induced responses. The directed migratory capacity of the G184S neutrophils to CXCL12 and CXCL2 declined as the neutrophils aged.

**Figure 5.**
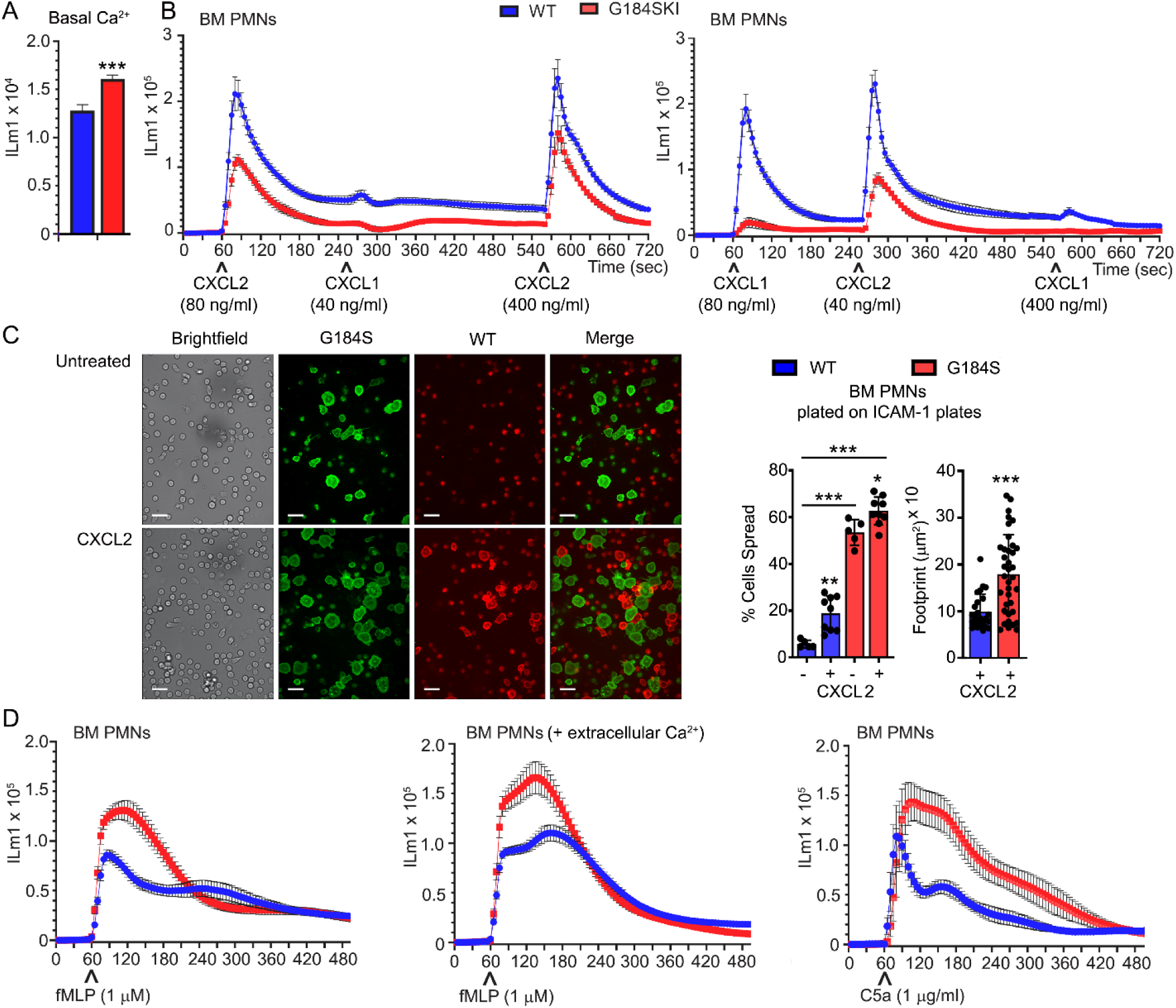
Assessing Chemokine Responsiveness and Integrin Mediated Adhesion of WT vs G184S Neutrophils *In Vitro*. (**A**) WT and G184S BM neutrophil basal fluorescence from 5 matched pairs of mice is shown. Statistics: data are means ± SEMs, then analyzed using unpaired Student’s *t*-test comparing G184S with WT. (**B**) Changes in intracellular calcium levels were monitored over 12 minutes (720 sec) in WT and G184S BM neutrophils exposed to CXCL2 and CXCL1, then re-stimulation with a higher dose of CXCL2 (left). Intracellular calcium response was again monitored over 12 minutes in WT and G184S BM neutrophils exposed to CXCL1, CXCL2, then re-stimulation with a higher dose of CXCL1 (right). Results are means ± SEMs from duplicate determinations and representative of 1 of 3 separate experiments. (**C**) Representative brightfield and confocal images showing CXCL2 triggered WT vs G184S BM neutrophil adhesion *in vitro* (right). Differentially labeled WT (red) and G184S (green) neutrophils treated with CXCL2 (bottom panels) or untreated (top panels) were seeded onto ICAM-1 coated plates. Scale bars = 20 µm. Their % spread and footprint sizes were quantified (right). Statistics: data are means ± SEMs, then analyzed using ANOVA. Results are from n ≥ 3 experiments done in duplicates. (**D**) The intracellular calcium response of WT vs G184S BM neutrophils monitored over 8 minutes (480 sec) after stimulations with fMLP (left, middle) and C5a (right). Results are means ± SEMs from duplicate determinations and representative of 1 of 3 separate experiments. (**A, B, D**) Data shown as fluorescent counts (ILm1). All results are from using WT and G184S BM reconstituted mice. *p < 0.05, **p < 0.005 and ***p < 0.0005.

### G184S Neutrophils Exhibit Defective *in vivo* Recruitment and TEM

To visualize how these *in vitro* defects translated *in vivo* we used the intravital microscopy to assess neutrophil behavior in the cremaster muscle blood vessels of a male mouse. In the initial experiment we injected IL-1β into the surrounding tissues and labeled the vasculature and neutrophils with a fluorescent CD31 and Gr-1 antibodies, respectively (Woodfin et al., 2011). Approximately 3 hours after IL-1β injection we imaged using mice either reconstituted with WT or G184S BM. Shown are snapshots taken at the beginning and 20 minutes after imaging commenced. Numerous WT neutrophils have been recruited and transmigrated, localized in the interstitium while significantly fewer G184S neutrophils had arrived or transmigrated (Figure 6A). Because of the poor release of G184S neutrophils into the bloodstream, we repeated the experiment, but in addition to local IL-1β, we injected AMD3100 intraperitoneally (Figure 6B). The addition of AMD3100 released more WT and G184S neutrophils into the circulation. This led to a marked increase in WT neutrophils in the cremaster blood vessels and massive amounts of transmigration (Video S3). Many more G184S neutrophils arrived, but relatively few transmigrated into the interstitium (Video S3). The massive TEM observed in WT mice was mostly due to efficient crossing of WT neutrophils through the endothelium after initial adherence, and their effective ability in finding appropriate transmigration sites (Video S4). In contrast, many G184S neutrophils engaged the endothelium, but failed to remain firmly attached, flying away or sliding along the endothelium without firmly adhering (Video S5). In addition, the few G184S neutrophils that underwent TEM had either spent significant longer time breaching the endothelial barrier or showed extensive intraluminal crawling before TEM (Video S6). We tracked 18 G184S neutrophil in a vessel segment over a 20-minute period after 90 minutes of IL-1β and AMD3100 treatment (Figure 6C). Five minutes after the initial image, 12 cells had departed the imaging field. Over the ensuing 15 minutes 1 more left and 2 transmigrated. One cell slid along the endothelium for 10 minutes before departing (Figure 6D). A comparison of the WT and G184S neutrophils shows many streaking G184S cells compared to WT neutrophils and fewer transmigration events both with IL-1β alone and with the addition of AMD3100 (Figure 6E). In order to evaluate neutrophil dynamics, we divided the tracked cells into two groups (attached to the vasculature and free) (Figure S3). A comparison of tracked cells within the blood vessels (attached cells) showed an increased speed and straightness of the G184S PMN tracks (Figure 6F). We also tracked neutrophils outside of the cremaster blood vessels and migrating within the interstitium (free cells). Representative tracks are shown for both WT and G184S interstitial neutrophils (Figure S4A). Once the G184S PMNs transmigrated they moved faster resulting in longer tracks and greater displacement (Figure S4B). Visualizing individual cells showed that the WT cells tended to adopt a polarized morphology with Gr-1 immunostaining concentrated on the uropod. In contrast, many of the G184S PMNs failed to properly polarize often exhibiting multiple lamellipodia and no clear uropod. Representative cells are shown (Figure S4C). Analyzing the movements of individual cells revealed the distorted movement of the leading and trailing edges (Figure S4D). Thus, the G184S neutrophils are poorly recruited to inflammatory sites often failing to adhere to the endothelium and undergo transmigration. Those few cells that transmigrate move quickly, but erratically.

**Figure 6.**
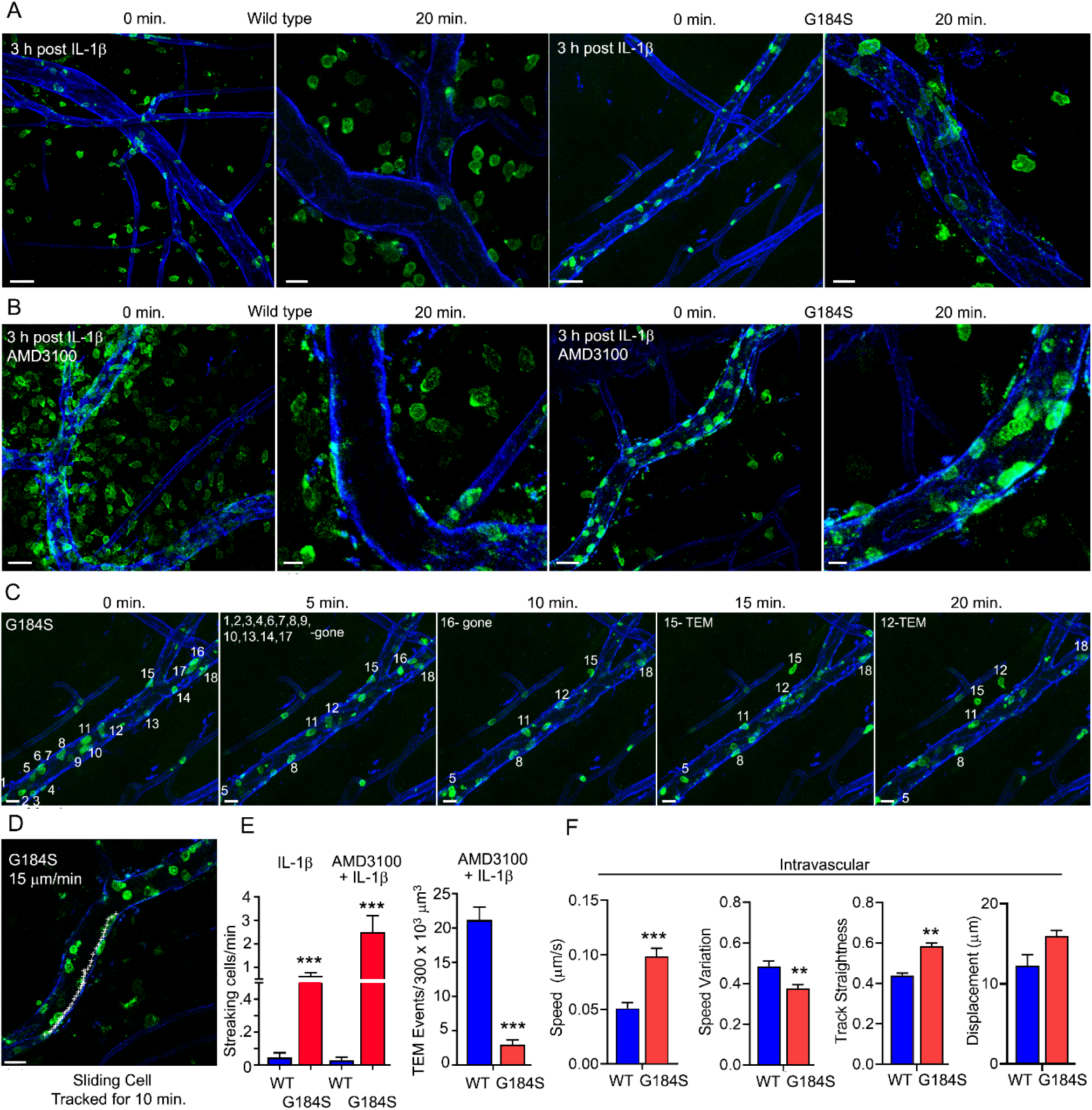
*In Vivo* TEM and Directed Migration of WT vs G184S Neutrophils during Acute Inflammation. (**A**) Representative confocal IVM images showing *in vivo* WT vs G184S neutrophil migration of WT and G184S BM reconstituted mice in the cremaster muscle subjected to 3 hrs of IL-1β (i.s.) treatment. Images Gr-1 immunostained neutrophils (green) at 0 and 20 min after IL-1β treatment are shown. Cremaster vasculatures (blue) are outlined by CD31 immunostaining. 0 min, scale bar = 40 µm; 20 min, scale bar = 15 µm. (**B**) Representative confocal IVM images showing *in vivo* WT vs G184S neutrophil migration in 3 hrs IL-1β (i.s.) and AMD3100 (i.p.) treated cremaster muscle. Images of 0 and 20 min after IL-1β and AMD3100 treatment are shown. Images are representative of WT (n = 3) and G184S (n = 3) BM reconstituted mice. 0 min, scale bar = 30 µm; 20 min, scale bar = 10 µm. (**C**) Representative confocal IVM images showing *in vivo* G184S neutrophil intravascular activities after 90 minutes of IL-1β (i.s.) and AMD3100 (i.p.) treated cremaster muscle. 18 G184S neutrophils were tracked over the course of 20 minutes with each neutrophil assigned to a number (white) at 0 min. Images at 0, 5, 10, 15, and 12 minutes are shown. Scale bar = 20 µm. (**D**) A representative confocal IVM image shows tracking (white +) of an intravascular G184S neutrophil crawling along the vessel lumen over 10 minutes. Scale bar = 20 µm. (**E**) Quantification of streaking cells over a 250 µm vessel segment in WT vs G184S mice treated with either IL-1β alone or IL-1β and AMD3100 (left). Transmigrated cells outside the vessel segment in WT vs G184S mice treated with IL-1β and AMD3100 (right). (**F**) Speed, speed variation, track straightness, and displacement from tracking intravascular WT vs G184S neutrophils in mice treated with IL-1β and AMD3100 over 30 minutes. (**A, E**) Results for only IL-1β treatment are from WT (n = 3) and G184S (n = 3) BM reconstituted mice for each study. (**C, D, E, F**) Results for 90 min treatment of IL-1β and AMD3100 are from WT (n = 5) and G184S (n = 5) BM reconstituted mice for each study. Statistics: data are means ± SEMs, then analyzed using unpaired Student’s *t*-test comparing G184S with WT. **p < 0.005 and ***p < 0.0005.

### Faulty G184S Neutrophils *In Vivo* Directed Migration

To test directed migration *in vivo*, we examined the recruitment of G184S neutrophils to a site of sterile injury. Recruitment to a sterile injury in the skin involves CXCR2, FPR2, and LTB4R1(Lammermann et al., 2013). It occurs in three phases; an initial Gα_i_-dependent scouting phase, a secondary amplification phase that likely depends upon the release of nicotinamide dinucleotide (NAD^+^) from damaged cells, and a stabilization phase (Ng et al., 2011). We imaged neutrophils in the lymph node following laser damage (LD). Using either WT or G184S BM reconstituted mice, we labeled the vasculature and neutrophils with intravenously injected fluorescent CD31 and Gr-1 antibodies, respectively (Figure S5A). Since the Gr-1 antibody labeled the intravascular neutrophils better than the lymph node resident neutrophils, we could distinguish the two-populations based on their fluorescence intensity. Within 5-10 minutes of the LD WT resident neutrophils appeared in the imaging field as they migrated towards the site. At approximately 20 minutes neutrophils increased in nearby blood vessels and within 30 minutes they had begun to transmigrate and move towards the LD site. By 45 minutes the transmigrated neutrophils had accumulated around the damage site (Video S6). In contrast, few resident or vascular G184S neutrophil arrived, and the majority of those that did arrive, failed to migrate towards the site of injury or to accumulate (Figure S5A, B and Video S6). Tracking individual neutrophils over the first hour shows the failure of G184S neutrophils to migrate towards the site of injury (Figure S5C) or to accumulate (Figure S5D). Both the transmigrated and residential neutrophils G184S neutrophils exhibited less directional migration and moved more erratically than did the WT neutrophils. The WT cell moved towards the site of LD while G184S cells often ignored the site and moved rapidly away (Figure S4E, F). The track parameters for the residential and transmigrated WT and G184S neutrophils are shown (Figure S5E). These results show that *in vivo* both tissue resident and transmigrated G184S neutrophils have a poor sense of direction consistent with a chemotaxis defect.

### G184S Neutrophils Fragment and Aggregate in Liver Sinusoids Following Concanavalin A Administration

The poor transmigration and aberrant aging of the G184S neutrophils prompted us to investigate whether the G184S neutrophils might cause or exacerbate vascular injury. The initial pathologic assessment of the lungs and liver in the WT and G184S BM reconstituted mice was unrevealing except for mild histologic evidence of hepatitis in both groups (data not shown). To assess the liver vasculature *in vivo* we used intravital microscopy. We injected fluorescently labeled Gr-1 and CD31 antibodies as previously. PBS injected G184S BM reconstituted mice had more neutrophils in their liver sinusoids, but otherwise did not differ from the controls (Figure 7A). Next, we treated the mice with Concanavalin A (ConA), which triggers inflammatory changes in the liver vascular bed (Siegmund et al., 2013). ConA can also trigger neutrophil extracellular trap formation, a potential contributing factor to vascular damage (Amulic et al., 2017). We administered a sublethal dose (2.5 mg/kg) intravenously and imaged 3-4 hours later. The G184S BM mice suffered pulmonary distress during imaging, while the WT BM reconstituted mice tolerated the procedure. Intravital microscopy revealed increased numbers of neutrophils in the liver sinusoids of the WT and G184S BM reconstituted mice, but more so in G184S BM reconstituted mice (Figure 7A and Video S7). Furthermore, more neutrophil fragments and aggregates were present in the G184S BM reconstituted mice liver sinusoids. Since neutrophils scan for activated platelets to initiate inflammation, we also imaged platelets in liver sinusoids of the ConA treated mice, which revealed platelet clots and aggregates, more prevalent in the G184S BM reconstituted mice (Figure 7B and Video S8). The ConA injection marginally elevated the WT and the G184S blood neutrophils (data not shown). Repeating the pathologic analysis following the ConA treatment revealed that the G184S livers had more liver sinusoid neutrophils and evidence of perivasculitis while similarly treated WT mice lacked the perivasculitis (Figure S6A). Perhaps explaining the pulmonary distress, slides prepared from the ConA treated G184S BM reconstituted mouse lungs revealed numerous neutrophil in the lung vasculature along with evidence of ongoing vascular inflammation, not observed in the controls (Figure S6B).

**Figure 7.**
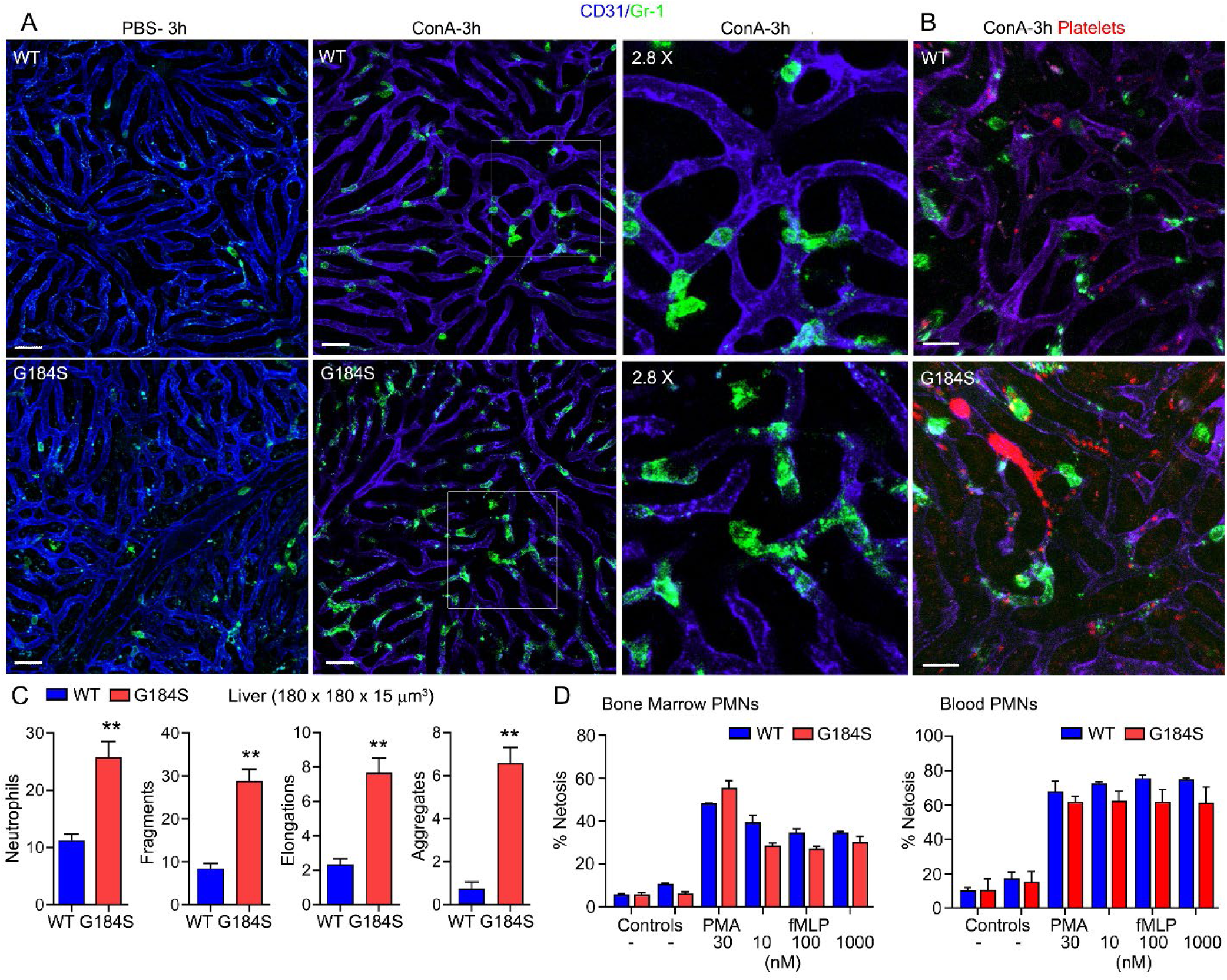
*In Vivo* Migration, Shape Change, Vitality, Platelet Interactions and *Ex Vivo* NET Formation of WT vs G184S Neutrophils during ConA-Induced Hepatitis. (**A**) Representative confocal IVM images of *in vivo* WT (top panels) vs G184S (bottom panels) neutrophil behaviors/characteristics in the liver after 3 hours of ConA (i.v.) treatment. Images of PBS-(i.v.) treated control mice (right) compared to ConA-treated mice (middle) shows Gr-1 immunostained neutrophils (green) migrating within the CD31 immunostained liver sinusoids (blue). Areas outlined by white squares are shown magnified in the right panels. Scale bar = 30 µm. (**B**) Representative confocal IVM images of *in vivo* WT (top) vs G184S (bottom) platelet – neutrophil interactions and platelet behaviors/characteristics in the liver after 3 hours of ConA treatment. Both Gr-1 stained neutrophils (green) and GP1bβ labeled platelets (red) are observed trafficking within the CD31 stained liver sinusoids (purple). Scale bar = 20 µm. (**C**) Quantifications of cell numbers, fragments, elongations, and aggregations within a 180 x 180 x 15 µm^3^ section for WT vs G184S neutrophils are shown. (**D**) Quantifications of ex vivo NET-formation assays from isolated WT vs G184S BM (left graph) and blood (right graph) neutrophils stimulated with PMA or increasing doses of fMLP. Results are from n = 3 separate experiments. (**A, B, C**) Results are from WT (n = 3) and G184S (n = 3) BM reconstituted mice for each study. Statistics: data are means ± SEMs, then analyzed using unpaired Student’s *t*-test comparing G184S with WT. **p < 0.005.

The increase neutrophil fragmentation *in vivo* suggested that the G184S neutrophils may have an increased propensity to form extracellular traps (Video S9). To test this possibility BM and blood neutrophils isolated from BM chimera mice were investigated to assess their propensity to form extracellular traps spontaneously or following treatment with phorbol myristate acetate (PMA) or with fMLP. Both PMA and fMLP trigger extracellular trap formation by stimulating NADPH oxidase to form reactive oxygen species. However, these assays did not reveal any intrinsic difference between the WT and G184S blood and bone marrow neutrophils (Figure 7D).

## Discussion

The G184S point mutation in the Gα_i2_ coding region disrupts the interaction between Gα_i2_ and RGS proteins. The failure of RGS proteins to bind Gα_i2_ slows the rate at which Gα_i2_-GTP returns to a GDP bound state. This has several consequences. First, the normal suppression of basal signaling provided by RGS proteins does not occur. Second, following GPCR triggered nucleotide exchange Gα_i2_-GTP can engage downstream effectors for longer durations. Furthermore, Gα_i2_-GTP does not recombine with freed G_βγ_ subunits, allowing them to persistently activate their own downstream effectors. This prolonged effector engagement by Gα_i_-GTP and free G_βγ_ subunits typically leads to hyperactivation of downstream signaling pathways (Kehrl, 2016; Woodard et al., 2015). Third, following ligand exposure the effective concentration of heterotrimeric Gα_i2_ available for GPCR activation declines. This can adversely affect the kinetics of the signal transduction pathways (Hwang et al., 2017). Cells including neutrophils that lack an individual RGS protein typically exhibit enhanced GPCR signaling (Balenga et al., 2014; Chan et al., 2018; Kehrl, 2016; Xie et al., 2016). Furthermore, studies of the G184S mice have largely reported augmented GPCR-linked Gα_i_ signaling (Neubig, 2015), yet past studies examining leukocyte chemoattractant receptors have largely reported diminished signaling (Cho et al., 2012; Hwang et al., 2015; Hwang et al., 2017). In the present study, we found that neutrophils, which carry the Gα_i2_ G184S mutation exhibit a complex phenotype with features of both hyperactive and hypoactive Gα_i_ signaling, which overlap with the phenotypes noted with Gα_i_ gain- or a loss-of function mutations.

A gain-of function mutation in a heterotrimeric Gα subunit typically results from the loss or diminution of its intrinsic GTPase activity. The Gα subunits remain persistently GTP bound unable to combine with Gβγ subunits, allowing both to engage downstream effectors. For example, a rat Gα_i2_ protein with a T182A mutation releases GDP more rapidly and hydrolyzes GTP more slowly than does the wild type protein, thereby prolonging its GTP bound status (Nishina et al., 1995). While the nearby G184S mutation does not affect GDP release, nor change the intrinsic Gα_i_ GTP hydrolysis rate; like the T182A mutation, it increases the duration that Gα_i2_ remains GTP bound. Consistent with an inappropriate or persistent engagement of downstream effectors, the G184S neutrophils have elevated basal intracellular calcium levels, increased basal cell motility, increased spreading on ICAM-1 coated plates, decreased basal pEzrin levels, and an altered cell morphology. These base line changes likely reflect the failure of RGS proteins to suppress ligand-independent, and low-level, spurious ligand-dependent GPCR signaling. A major functional role for neutrophil RGS proteins may be to suppress background noise in chemoattractant receptor signaling pathways, thereby keeping the cell poised to respond to a strong chemoattractant signal. A similar role for RGS protein/Gα_i2_ interactions has been suggested in platelets, where they help provide a threshold for platelet activation (Signarvic et al., 2010). Compared to a Gα_i2_ G184S mutation, a T182A mutation would more severely disrupt GPCR-mediated Gα_i2_ signaling and likely result in a more adverse neutrophil phenotype. In fact, humans carrying a Gα_i2_ T182 mutation on one allele have innate immune defects and suffer recurrent sinopulmonary infections likely explained by neutrophil defects (Lamborn et al., 2016). No humans with a G184S Gα_i_ mutation have been reported. A G184S mutation on one allele may have a relatively mild phenotype, but homozygotes would likely have a severe multisystem phenotype. Similarly, a mouse or human carrying two Gα_i2_ T182A alleles would likely not be viable.

A loss-of-function of mutation in Gα_i2_ such as G203T sequesters Gβγ subunits and GPCRs in their inactive conformations (Inoue et al., 1995; Murray-Whelan et al., 1995). While no such mutation in Gα_i2_ has been reported in humans or engineered in mice, neutrophils with such a mutation would be expected to respond poorly to chemoattractants, resembling neutrophils that lack specific receptors or critical elements in the signaling pathway such as Gα_i_ (Kehrl, 2016). Neutrophils lacking Gα_i2_ exhibit impaired interstitial chemotaxis and poor accumulation at laser damage sites (Lammermann et al., 2013). A G184S mutation can mimic a loss-of-function mutation by limiting heterotrimer availability, by reducing GPCR/G-protein coupling, and by decreasing receptor availability. In neutrophils the G184S mutation caused a progressive loss of CXCR2 expression and CXCL1/2 triggered chemotaxis. Thus, the excessive engagement of effectors in the Gα_i2_ signaling pathway as neutrophils develop and age causes a progressive decline in CXCR2 expression and signaling. This was already evident even in bone marrow neutrophils, which fail to mobilize properly to CXCL2. In contrast, CXCR4 expression in G184S neutrophils resembled that of WT cells and CXCR4 signaling in bone marrow neutrophils remained largely intact even exaggerated at low ligand concentrations. However, eventually CXCL12 mediated chemotaxis declined and splenic neutrophil migrated poorly despite their adequate receptor expression. Thus, CXCR4 expression is not sensitive to hyperactive Gα_i2_ signaling, but eventually signaling is affected. The reason for this progressive loss is unclear. Perhaps the inability to suppress low grade signals over time downregulates critical effectors in the chemotaxis signaling pathway. Further evidence of poor CXCR2 signaling in peripheral G184S neutrophils was their poor adherence to vascular endothelium *in vivo* and their inability cross endothelial borders. This is despite their increased propensity to spread on ICAM-1 coated plates *in vitro* and it underscores the necessity of finely regulated Gα_i2_ mediated integrin activation for neutrophil adhesion to endothelial cells under flow for eventual neutrophil transmigration. Additional evidence of poor chemoattract receptor function, the G184S neutrophil that managed to transmigrate exhibit difficulties in maintaining a leading edge and following endogenous chemoattractant gradients.

The severe defects in neutrophil trafficking arise from the impact of the G184S Gα_i2_ mutation on chemoattractant receptor signaling pathways, however, less clear are the causes of the neutrophil fragmentation and vascular inflammation noted after treating the G184S BM reconstituted mice with ConA. We found no intrinsic difference in BM and blood neutrophil extracellular trap formation in response to PMA or to fMLP arguing that that an increased propensity to form extracellular traps likely does not explain the ongoing vascular inflammation. Another possibility is abnormal neutrophil aging in these mice. Normal neutrophil aging begins after neutrophils leave the BM. Eventually aged neutrophils migrate into healthy tissues, a process termed clearance. This helps protects the vascular system from neutrophil mediated thrombo-inflammation (Adrover et al., 2019; Casanova-Acebes et al., 2018). Due to their severe TEM defect, the G184S mice neutrophils accumulate in both the lungs and liver vasculature. The TEM defect interferes both with the recruitment of neutrophils to inflammatory sites, but also with normal neutrophil clearance, thereby explaining the neutrophil overload. The persistent presence of aged neutrophils in the lung and liver blood vessels in the setting of an inflammatory insult may damage blood vessels and contributed to the ongoing thrombo-inflammation noted in the G184S mice following ConA treatment. However, we cannot exclude a role for abnormal platelet function as a contributing factor since the platelets in the G184S BM reconstituted mice also carry the G184S Gα_i2_ alleles (Signarvic et al., 2010).

Our studies underscore two essential functions for RGS proteins in neutrophils. First, they suppress low grade environmental signals maintaining a low basal level of activity in Gα_i_-linked signaling pathways. This keeps the cells ready and able to respond to chemoattractant signals. Second, they help coordinate both the magnitude and duration of signals generated through Gα_i_ linked signaling. Interrupting these essential functions in neutrophils disrupts neutrophil homeostasis, trafficking, and function. Pharmaceutically targeting all Gα_i_/RGS interactions is likely to severely disable neutrophils. A more finely tuned intervention will require a better understanding of how individual RGS proteins cooperate to regulate chemoattractant receptor signaling.

## Material and methods

### Key resources table

| REAGENT or RESOURCE | SOURCE | IDENTIFIER |
| --- | --- | --- |
| **Experimental Models:**<br>**Organisms/Strains** |  |  |
| Mouse, C57BL/6J | The Jackson Laboratory | JAX 000664 |
| Mouse, B6.129S1-*Gnai2*<sup>tm1.1Rneu</sup>/J | The Jackson Laboratory | JAX 012768 |
| Mouse, B6.SJL-Ptprc<sup>a</sup> Pepc<sup>b</sup>/BoyJ | The Jackson Laboratory | JAX 002014 |
| Mouse, LysM-eGFP | Dr. Thomas Graf | N/A |
| **Antibodies** |  |  |
| Anti-mouse CD11b-APC-Cy7 (clone M1/70) | BD Pharmingen | Cat#553312; RRID: AB_396772 |
| Anti-mouse CD11c-Biotin (clone HL3) | BD Pharmingen | Cat#553800; RRID: AB_395059 |
| Anti-mouse F4/80-Biotin (clone BM8) | BioLegend | Cat#123106; RRID: AB_893501 |
| Anti-mouse Gr-1-Biotin (clone RB6-8C5) | BD Pharmingen | Cat#1088430; RRID: AB_394641 |
| Anti-mouse Ly6C-V450 (clone AL-21) | BD Pharmingen | Cat#560453; RRID: AB_1645284 |
| Anti-mouse Ly6G-PE Cy7 (clone 1A8) | BD Pharmingen | Cat#560601; RRID: AB_1727562 |
| Anti-mouse NK1.1-Biotin (clone PK136) | BD Pharmingen | Cat# 553163; RRID: AB_394675 |
| Anti-mouse TCR$\gamma\delta$-Biotin (clone GL3) | BD Pharmingen | Cat# 553176; RRID: AB_394687 |
| Anti-mouse B220-Biotin (clone RA3-6B2) | BD Pharmingen | Cat#553086; RRID: AB_394616 |
| Anti-mouse CD117-Biotin (clone 1B8) | BD Pharmingen | Cat# 553353; RRID: AB_394804 |
| Anti-mouse TER-119-Biotin (clone TER-119) | BD Pharmingen | Cat#553672; RRID: AB_394985 |
| Anti-mouse CD3-Biotin (clone 145-2C11) | BioLegend | Cat#100304; RRID: AB_312669 |
| Anti-mouse CD4-Biotin (clone GK1.5) | BioLegend | Cat#100404; RRID: AB_312689 |
| Anti-mouse CD5-PerCP Cy5.5 (clone 53-7.3) | BioLegend | Cat#100624; RRID: AB_2563433 |
| Anti-mouse CD8-Biotin (clone 53-6.7) | BioLegend | Cat#100704; RRID: AB_312743 |
| Anti-mouse CD62L-PE (clone Mel-16) | BioLegend | Cat#104408; RRID: AB_313095 |
| Anti-mouse CD45.1-FITC (clone A20) | BioLegend | Cat#110706; RRID: AB_313495 |
| Anti-mouse CD45.2-APC (clone 104) | BioLegend | Cat#109814; RRID: AB_389211 |
| Anti-mouse CD182-PE (CXCR2, clone SA045E1) | BioLegend | Cat#149610; RRID: AB_2565690 |
| Anti-mouse CD184-PE (CXCR4, clone 2B11) | Thermo Fisher Scientific | Cat#12-9991-82; RRID: AB_891391 |
| Anti-mouse CD169-A488 (clone 3D6.112) | BioLegend | Cat#142419; RRID: AB_2566436 |
| Anti-mouse Ly6G-A594 (clone 1A8) | BioLegend | Cat#127636; RRID: AB_2563207 |
| Anti-mouse CD31-APC (clone 390) | eBioscience | Cat#17-0311-82; RRID: AB_657735 |
| Anti-mouse Ly6G-FITC (clone 1A8) | eBioscience | Cat#11-9668-82; RRID: AB_2572532 |
| Anti-mouse CD45-BV650 (clone 30-F11) | BioLegend | Cat#103151; RRID: AB_2565884 |
| Anti-mouse CD31, purified (clone 390) | BioLegend | Cat#102402; RRID: AB_312897 |
| Anti-mouse Gr-1, purified (clone RB6-8C5) | BioLegend | Cat#108402; RRID: AB_313367 |
| Anti-GPIIbβ-DyLight649 (derivatized) | Emfret Analytics | Cat#X649; RRID: AB_2861336 |
| Streptavidin-BV605 | BioLegend | Cat#4052259 |
| **Chemicals, Peptides, and Recombinant Proteins** |  |  |
| Sphingosine-1-phosphate (d18:1) | Cayman | Cat#62570-1MG |
| fMLP (N-Formyl-Met-Leu-Phe) | Tocris | Cat#1921-1MG |
| Recombinant murine Complement Component C5a | R&D Systems | Cat#2150-C5-025 |
| Recombinant murine CXCL1 (KC) | R&D Systems | Cat#453-KC |
| Recombinant murine CXCL2 | PeptoTech | Cat#250-15 |
| Recombinant murine CXCL12 | PeptoTech | Cat#250-20A |
| Bovine Serum albumin fatty acid free (low endotoxin) | Sigma-Aldrich | Cat#8806 |
| AMD 3100 | Sigma-Aldrich | Cat# A5602-5MG |
| KRH 3955 | Tocris | Cat#6126-10MG |
| PMA | Sigma-Aldrich | Cat#P8139 |
| Helix NP<sup>TM</sup> NIR | BioLegend | Cat#425301 |
| DAPI | BioLegend | Cat#422801 |
| Avertin (Tribromoethanol) | Sigma-Aldrich | Cat#T48402-5G |
| Evans Blue Dye | Sigma-Aldrich | Cat#E2129 |
| Recombinant murine ICAM-1-FC | R&D Systems | Cat#796-IC-050 |
| Recombinant murine IL-1β | R&D Systems | Cat#401-ML-005 |
| Concanavalin A (ConA) | InvivoGen | Cat#inh-cona |
| **Critical Commercial Assays** |  |  |
| Alexa Fluor 488 protein labeling kit | Thermo Fisher Scientific | Cat#A10235 |
| Alexa Fluor 555 protein labeling kit | Thermo Fisher Scientific | Cat#A20174 |
| Alexa Fluor 568 protein labeling kit | Thermo Fisher Scientific | Cat#A10238 |
| Alexa Fluor 647 protein labeling kit | Thermo Fisher Scientific | Cat#A20173 |
| FLIPR Calcium 4 assay kit | Molecular Devices | Cat# R8142 |
| LIVE/DEAD™ Fixable aqua dead cell stain kit | Thermo Fisher Scientific | Cat#L34965 |
| LIVE/DEAD™ Fixable yellow dead cell stain kit | Thermo Fisher Scientific | Cat#L34967 |
| APC BrdU Flow kit | BD Biosciences | Cat#552598 |
| Anti-Ly-6G MicroBeads UltraPure, mouse | Miltenyi Biotec | Cat#130-120-337 |
| **Software and Algorithms** |  |  |
| SOFT max Pro 5.2 | Molecular Devices | https://www.moleculardevices.com/ |
| FlowJo v10.7 | Tree Star | https://www.flowjo.com/ |
| ImageJ 1.53a | NIH | https://imagej.nih.gov/ij/ |
| Imaris v9.3.1 | Bitplane | http://www.bitplane.com/Imaris |
| Prism v8.0.1 | GraphPad | https://www.graphpad.com/scientific-software/prism/ |

Resource Availability-Further information and requests for resources and reagents should be directed to and will be fulfilled by the Lead Contact, John H. Kehrl

Materials Availability-All animal strains used in this study are available from The Jackson Laboratory including the B6.129S1-*Gnai2*^*tm1*.*1Rneu*^/J and the B6.129P2-*Lyz2*^*tm1(cre)Ifo*^/J with exception of the LysM-eGFP mice, which are available with permission from Dr. Thomas Graf (Center for Genomic Regulation, Barcelona, Spain).

### Mice

Male (20-25g) mice were used. C57BL/6 and B6.SJL-Ptprc^a^ Pepc^b^/BoyJ mice were obtained from the Jackson Laboratory. B6.129S1-*Gnai2*^*tm1*.*1Rneu*^/J (G184S), Jackson Laboratory mice were backcrossed >17 times onto C57BL/6. Litter mate controls were used for experiments that directly compared wild-type (WT) and G184S mice. The G184S mice were bred as heterozygotes. All mice were maintained in specific-pathogen-free conditions at an Association for Assessment and Accreditation of Laboratory Animal Care-accredited animal facility at the NIAID. All procedures and protocols regarding animal experiments were performed under a study protocol approved by the NIAID Animal Care and Use Committee (National Institutes of Health). For the BM reconstitutions 6 wk-old C57BL/6 (CD45.1) mice were irradiated twice with 550 rad for total of 1100 rad and received BM from C57BL/6 CD45.2 mice (control) or from G184S CD45.2 mice. Mixed chimeric mice were made by reconstituting C57BL/6 CD45.1 mice with a 1:1 mix of BM from C57BL/6 CD45.2 mice (WT) and from G184S CD45.2 mice. The engraftment was monitored by sampling the blood 28 d later. The mice were used 6 – 7 wk after reconstitution. All mice used in this study were 6 – 12 weeks of age.

### Cell isolation

BM cells were isolated from mouse femur and tibia. BM and splenic neutrophils were purified to a purity of ∼95% using anti-Ly-6G MicroBeads UltraPure (Miltenyi Biotech). When needed neutrophils were cultured in RPMI 1640 containing 10% fetal calf serum (FCS, Gibco), 2 mM L-glutamine, antibiotics (100 IU/mL penicillin and 100 μg/mL streptomycin), 1 mM sodium pyruvate, and 50 µM 2-mercaptoethanol. Cell culture media for S1P chemotaxis was same as above except charcoal-dextran filtered FCS serum was used.

### Flow cytometry

Single cells were re-suspended in PBS, 2% FCS, and stained with fluorochrome-conjugated or biotinylated antibodies against CD11b (M1/70), Ly6G (1A8), Ly6C (AL-21), CD11c (HL3), F4/80 (BM8), NK1.1 (PK136), TCRγδ (GL3), B220 (RA3-6B2), CD117 (2B8), TER-119 (TER-119), CD3 (145-2C11), CD4 (GK1.5), CD5 (53-7.3), CD8 (53-6.7), CD184 (CXCR4, 2B11), CXCR2 (SA044E1), CD62L (MEL-16), CD45.1 (A20), or CD45.2 (104) (all from Thermo Fisher, BioLegend, or BD Pharmingen). Biotin-labeled antibodies were visualized with fluorochrome-conjugated streptavidin (Thermo Fisher, or BioLegend). LIVE/DEAD™ Fixable Aqua Dead Cell Stain Kit, or LIVE/DEAD™ Fixable Yellow Dead Cell Stain Kit (Thermo Fisher) were used in all experiments to exclude dead cells. Data acquisition was done on FACSCanto II and FACSCelesta SORP (BD) flow cytometer and analyzed with FlowJo software (Treestar). For intracellular flow cytometry cells were intracellular stained using the eBioscience™ Intracellular fixation & permeabilization buffer Set (Thermo Fisher) protocol using the PE conjugated CXCR2, PE-conjugated CXCR4 antibody, phallodin, or pERM antibody. To detect phosphorylated ERM proteins a rabbit anti-phospho–ezrin (Thr^567^)/radixin (Thr^564^)/moesin (Thr^558^) (pERM) antibody was used (Cell Signaling Technology). Isotype control staining was performed using rabbit IgG isotype mAb Alexa Fluor 647 (DA1E; Cell Signaling Technology). Secondary F(ab′)_2_ fragment of goat anti-rabbit IgG (H+L) Alexa Fluor 647 (Thermo Fisher Scientific) was used to detect the pERM Abs. To detect F-actin the cells were stained using Alexa Fluor 568 or 647 conjugated to phalloidin (Invitrogen). After washing, the cells were resuspended in 250 μl of 1% BSA/PBS and filtered prior to acquisition on the flow cytometer.

### BrdU labeling

BrdU labeling of endogenous neutrophils was modified from previously described (Uhl et al., 2016). Briefly, neutrophil precursors in the mouse BM were labeled via intravenous (i.v.) injection of 5-bromo-2′-deoxyuridine (BrdU; 2 mg per mouse; FITC (fluorescein isothiocyanate) BrdU Flow Kit; BD Biosciences. 2 days after BrdU injection, mice received i.v. injection of 100 μg CD62L antibody to block transendothelial migration and to reduce neutrophil margination.

### Neutrophil mobilization

Neutrophil mobilization to the blood was performed as previously described (Martin et al., 2003). Mice were injected with CXCL1 (40 μg/kg), AMD 3100 (5mg/kg), KRH 3955 (2.5mg/kg), or PBS only via the tail vein. A group of mice received intravenously both AMD3100 and CXCL1. The AMD 3100 was injected 20 min prior to the CXCL1. Blood was collected before and 1, 2, and 3 hours after injection from the mandibular plexus or the tail vein. The samples were analyzed by flow cytometry.

### Chemotaxis and migration assays

Chemotaxis assays were performed using a transwell chamber (Costar) as previously described (Hwang et al., 2017). BM cells or splenocytes were immunostained for neutrophils with fluorochrome-conjugated antibodies against CD11b, Ly6C, and Ly6G, washed twice, re-suspended in complete RPMI 1640 medium and added in a volume of 100 μl to the upper wells of a 24-well transwell plate with a 5 µm insert. Lower wells contained various doses of chemokines in 600 μl of complete RPMI 1640 medium. The numbers of cells that migrated to the lower well after 1 hr incubation were counted using a MACSQuant flow cytometer (Miltenyi Biotec). The percent migration was calculated by the numbers of neutrophils that migrated into the bottom chamber divided by the total number of neutrophils in the starting cell suspension and multiplying the results by 100. D-erythro-sphingosine 1-phosphate was purchased from Sigma. CXCL1, fMLP, and C5a used for the migration assays were purchased (R&D Systems) CXCL2, and CXCL12 were purchased (PeproTech). Fatty acid free bovine serum albumin (FAF-BSA) was purchased (Sigma-Aldrich).

### Intracellular calcium measurement

Cells were seeded at 10^5^ cells per 100 µl loading medium (RPMI 1640, 10% FCS) into poly-D-lysine coated 96-well black wall, clear-bottom microtiter plates (Nalgene Nunc). An equal volume of assay loading buffer (FLIPR Calcium 4 assay kit, Molecular Devices) in Hank’s balanced salt solution supplemented with 20 mM HEPES and 2 mM probenecid was added. Cells were incubated for 1 hr at 37°C before adding the indicated concentration of chemokine, fMLP or C5a and then the calcium flux was measured using a FlexStation 3 (Molecular Devices). The data was analyzed with SOFT max Pro 5.2 (Molecular Devices). Data is shown as fluorescent counts and the y-axis labeled as Lm1.

### Receptor internalization

BM cells or purified blood neutrophils were rested in RPMI 1640/10 mM HEPES for 30 min at 37°C/5% CO_2_ before immunostaining for CXCR2. Subsequently the cells were stimulated with fMLP (1 μM), solvent (DMSO), CXCL1 (100 ng/ml), or CXCL2 (100 ng/ml), while being maintained at 37°C/5% CO_2_. Thirty minutes later the cells were fixed and analyzed by flow cytometry for cell surface CXCR2 expression.

### Adhesion assay

Glass-bottom culture dishes (MatTek) were coated with 1 μg/ml ICAM-1 in phosphate-buffered saline (PBS) containing 2 mM MgCl_2_ and 1 mM CaCl_2_ (200 μl/well) at 4°C overnight. The dishes were washed three times with PBS before use. Neutrophils were isolated from BM cells by positive selection using the neutrophil isolation kit (Miltenyi Biotec, Auburn, CA). WT and G184S KI neutrophils were stained by Alexa Fluor 568- and Alexa Fluor 488-conjugated anti-mouse Gr-1 antibodies, respectively. Cells were rested at 37°C for 30 min. Equal numbers of WT and G184S KI neutrophils (1 x 10^5^ cells/100 ul) were transferred to a single tube to which was added 100 ng/ml CXCL2 and 5 mM MgCl_2_. Cells were immediately seeded on a ICAM-1 coated dish and placed in an incubator at 37°C and 5% CO_2_ for 30 min. Images were acquired on a PerkinElmer UltraVIEW spinning disc confocal system (PerkinElmer Life Science, Waltham, MA), with a Zeiss Axiovert 200 inverted microscope (Carl Zeiss Microimaging, Thornwood, NY) equipped with a 40X oil-immersion objective (N.A. 1.3). Images were acquired and the data were processed using ImageJ software (National Institutes of Health, Bethesda, MD).

### Analysis of neutrophil aging in blood

Analysis of neutrophil aging was performed as previously described (Adrover et al., 2019). Briefly, we compared blood drawn at ZT5 or ZT13 in mice housed under a 12h:12h light:dark cycles. We collected blood at the indicated time points and analyzed blood neutrophil numbers and their expression levels of CXCR4, CXCR2, or CD62L. Blood neutrophils at ZT5 were also analyzed for F-actin levels and pERM levels by intracellular flow cytometry as describe above. Forward light scatter of ZT5 and ZT13 neutrophils was collected by flow cytometry. Previously published RNA-sequencing data (Genome Expression Omnibus accession number GSE86619) was used to determine RGS protein and Gα protein mRNA expression in ZT5 and ZT13 mouse neutrophils(Adrover et al., 2019).

### Neutrophil NETosis

BM Neutrophils (> 95% purity) were resuspended in RPMI medium containing 10% FCS (1 x 10^6^/ml) and incubated for 37°C and 5% CO_2_ for 30 min. To induce NETosis, the cells were exposed to the activating agent PMA (10, 30 nM; Sigma-Aldrich) or fMLP (100, 1000 nM; Sigma-Aldrich) for 0-4 hrs at 37°C. NET induction was terminated by 4% PFA for 15 min. Helix NP™ NIR (0.1 µM; Biolegend) and DAPI (0.3 nM; Biolegend) were added to detect NETs. Data acquisition (Helix NP™ NIR^+^ DAPI^+^ Cells) was done on FACSCelesta SORP (BD) flow cytometer and analyzed with FlowJo software (Treestar).

### Spleen imaging

Immunohistochemistry was performed as previously described (Park and Kehrl, 2019). Briefly, freshly isolated spleens were fixed in 4% paraformaldehyde (Electron Microscopy Science) overnight at 4°C on an agitation stage. Spleens were embedded in 4% low melting agarose (Thermo Fisher Scientific) in PBS and sectioned with a vibratome (Leica VT-1000 S) at a 30 µm thickness. Thick sections were blocked in PBS containing 10% fetal calf serum, 1 mg/ml anti-Fcγ receptor (BD Biosciences), and 0.1% Triton X-100 (Sigma-Aldrich) for 30 min at room temperature. Sections were stained at 4°C on an agitation stage overnight with anti-CD169 and anti-Ly6G antibodies (BioLegend). Stained sections were microscopically analyzed using Leica SP8 confocal microscope (Leica Microsystem) equipped with an HC PL APO CS2 40X oil-immersion objective NA, 1.3. Images were processed with Leica LAS AF software (Leica Microsystem) and Imaris software v.9.0.1 64x (Bitplane AG).

### BM imaging

For imaging neutrophil recruitments from the BM, mice were injected i.v. with CXCL1 (40 μg/kg) or with an intraperitoneal (i.p.) injection of AMD3100 (5 mg/kg). BM surgery preparation for intravital microscopy was modified from a previously described protocol (Lo Celso et al., 2011). Briefly, anesthesia was initiated with an i.p. injection of Avertin (300 mg/kg, tribromoethanol; Sigma-Aldrich). Blood vessels were outlined by the i.v. injection of 1% EB solution in PBS (Evans Blue Dye, Sigma-Aldrich) at 1 ml/kg. A single incision, starting between the ears and following the head midline until 3-4 mm from the nose area was made with sharp scissors, then skin flaps were separated by pulling toward the sides with forceps. After surgery, the mice were transferred and stabilized onto an imaging stage using an upright setup and a custom-made stage with a head holder (NIH Division of Scientific Equipment and Instrumentation Services). For anesthesia isoflurane (Baxter; 2% for induction, 1 – 1.5% for maintenance, vaporized in an 80:20 mixture of oxygen and air) was used. 2-photon laser scanning microscopy was performed with a LEICA SP5 inverted 5-channel confocal microscope (Leica Microsystems) equipped with a 25X water-immersion objective, N.A. 0.7. 2-photon excitation was provided by a Mai Tai Ti:Sapphire laser (Spectra Physics) with a 10 W pump, tuned wave length rages from 820 – 920 nm. Emitted fluorescence was collected using 4-channel non-descanned detector. Wavelength separation was through a dichroic mirror at 560 nm and then separated again through a dichroic mirror at 495 nm followed by 525/50 emission filter for GFP; 405/20 emission filter for second harmonics; and the Evans blue signal was collected by 680/50 nm emission filter. For 4D analysis of cell migration, stacks of 4 or 10 sections (z step = 5 µm) were acquire every 15 or 30 s to provide an imaging volume of 20 or 50 µm in depth. Post-acquisition mages were processed using Imaris (Bitplane) software.

### Lung imaging

Confocal microscopy of live lung sections was developed as a technique for visualizing tissue architecture, cell segregation and cell migration in mouse lung *in vitro*, at as close to physiological conditions as possible. After euthanasia, mouse lungs were inflated with 1.5 % of low-melt agarose in RPMI at 37 C. Inflated tissues were kept on ice, in 1 % FCS in PBS, and sliced into 300-350 µm sections using Leica VT1000 S Vibrating Blade Microtome (Leica Microsystems) at speed 5, in ice-cold PBS. Tissue sections were stained with fluorescently labeled anti-CD31, anti-CD45, and anti-Ly6G antibodies (eBioscience) for 2 h on ice. After staining sections were washed 3 times and cultured in complete lymphocyte medium (Phenol Red-free RPMI supplemented with 10 % FCS, 25 mM HEPES, 50 μM β-ME, 1 % Pen/Strep/L-Glu and 1 % Sodium Pyruvate) in humidified incubator at 37° C. Tissues were allowed to completely recover for 12 h prior to imaging. Sections were held down with tissue anchors (Warner Instruments) in 14 mm microwell dishes (MatTek), and imaged using Leica DMi8 inverted 5 channel confocal microscope equipped with an Environmental Chamber (NIH Division of Scientific Equipment and Instrumentation Services) to maintain 37° C and 5 % CO_2_. Microscope configuration was set up for four-dimensional analysis (x,y,z,t) of cell segregation and migration within tissue sections. Diode laser for 405 nm excitation; Argon laser for 488 and 514 nm excitation, DPSS laser for 561; and HeNe lasers for 594 and 633 nm excitation wavelengths were tuned to minimal power (between 1 and 5 %). Z stacks of images were collected (10 – 50 µm). Mosaic images of lung sections were generated by acquiring multiple Z stacks using motorized stage to cover the whole section area and assembled into a tiled image using LAS X (Leica Microsystems) software. For time-lapse analysis of cell migration, tiled Z-stacks were collected over time (1 to 4 h). Post-acquisition mages were processed using Imaris (Bitplane) software.

### Preparation of cremaster muscle for imaging

For imaging neutrophils in the cremaster muscle, an intrascrotal injection of IL-1β (50 ng in 300 µl saline; R&D Systems) and in some instances an i.p. injection of AMD3100 (5 mg/kg in 200 µl saline; R&D Systems) were used to stimulate acute inflammation in the cremaster muscle and to mobilize neutrophils. Separate saline injections were used as controls. After 90 min, the mice received injections of Avertin (300 mg/kg, i.p.) and fluorescently labeled antibodies (i.v.) to visualize neutrophils and blood vessel endothelium. Subsequently, the right cremaster tissue was exposed via a scrotal skin incision, and carefully freed by dissecting it away from the associated connective tissues. The isolated cremaster tissue was exteriorized and stabilized onto the imaging stage/insert with the tissue directly contacting the cover glass. The exposed tissue was kept moist with pre-warmed saline (37°C).

### Preparation of the inguinal lymph node for imaging

For imaging neutrophils in lymph nodes, mice received injections of Avertin (300 mg/kg, i.p.) and fluorescently labeled antibodies (i.v.) to delineate neutrophils and venule endothelium. Inguinal lymph nodes were prepared for intravital microscopy as previously described (Yan et al., 2019). Briefly, the inguinal lymph node was surgically exposed without perturbing blood and lymph vessels, stabilized on the imaging stage/insert directly contacting the cover glass, and kept moist with pre-warmed saline. To induce laser damage, the desired target area was set at the middle optical plane of the z-stack using a zoom factor of 25. The Mai Tai laser power was set to 50% for 3 sec.

### Preparations for liver imaging

To image liver neutrophils, the mice received an i.v. injection of saline or ConA (2.5 mg/kg, Sigma-Aldrich) to induce liver inflammation. Three hours later, the mice received Avertin (300 mg/kg, i.p.) and fluorescently labeled antibodies (i.v.) to outline liver sinusoids, neutrophils and platelets. To expose the liver an incision was made along the midline on the abdomen from top of the bladder area to the sternum was made. The skin on the left and right side from abdominal muscles were separated, then cut and removed. A second incision along the midline of the abdomen exposed the xiphoid cartilage, and the abdominal muscles were removed by cauterization. Mice were placed on an imaging stage/insert, and carefully stretched to allow the liver to be exposed. An exposed liver lobe was flipped onto the cover glass and covered with a saline-moistened gauze. Confocal intravital imaging was performed as described below.

### Imaging neutrophils in the cremaster, lymph node, and liver

Once stabilized onto an imaging stage/insert the mouse received isoflurane (Baxter; 2% for induction of anesthesia, and 1 – 1.5% for maintenance, vaporized in an 80:20 mixture of oxygen and air), and placed into a temperature-controlled chamber. A Leica SP8 inverted 5 channel confocal microscope equipped with a motorized stage, 4 hybrid ultra-sensitive detectors, and argon and helium neon lasers, and a 25x water-immersion objective (1.0 N.A) was used for imaging (Leica Microsystems). This microscope incorporates optical zoom function, and most image sequences were captured at a final magnification of x25-x75. For cremaster imaging, sequential scans of the 488-nm channels for anti-Gr-1 AlexaFluor-488 labeled neutrophils (10 µg/mouse; clone RB6-8C5) and 633-nm channels for anti-CD31-AlexaFluor-647 labelled unbranched 25-40 μm venules (10 µg/mouse; clone 390) were acquired. For the inguinal lymph nodes, sequential scans of the 488-nm channels for anti-Gr1 AlexaFluor-488 labeled neutrophils (10 µg/mouse) and 561-nm channels for anti-CD31-AlexaFluor-555 labeled HEVs (10 µg/mouse) were captured following laser damage. For the liver, scans of the 488-nm channels were used to detect anti-Gr1 AlexaFluor-488 labeled neutrophils (10 µg/mouse), 561-nm channels for anti-CD31-AlexaFluor-568 labeled sinusoids (10 µg/mouse), and 633-nm channels for anti-GPIbβ-DyLight649 labelled platelets (2µg/mouse) (Grosse et al., 2007). Images were acquired at a resolution of 1024 x 1024 pixels, which corresponds to a voxel size of approximately 0.25 x 0.25 x 0.7 µm in the x, y, z planes. Image stacks of optical sections ∼ 1 µm thickness were routinely acquired at 20 – 40 sec intervals, and with the incorporated resonance scanner of 8,000 Hz. This methodology allowed acquisition of full 3D confocal images of cremaster venules, HEVs/laser damage, and liver sinusoid, which yielded high-resolution 4D videos (real-time in 3D) of dynamic events. Tissues were firmly maintained for all preparations without (or with minimal) movement interfering with blood flow or image acquisition.

### Intravital image processing and analysis of leukocyte dynamics

After acquisition, sequences of z-stack images were analyzed with Imaris Software (version 9.5.0, Bitplane AG, Zurich, Switzerland. For cremaster confocal intravital imaging sequential z-sections of stained cells in time frames were acquired for 3D reconstruction and surface modeling of representative. The 3D cell surfaces were then tracked using tracking algorithm of Imaris. The tracked cells were then divided into two groups (attached to the vasculature and free) utilizing the filtering function of Imaris and mean intensity threshold of the channel representing veins.

### Histopathology

Mice were injected with ConA (2.5 mg/kg in in 90 µl saline, i.v.) for liver confocal intravital imaging experiments. Mice were sacrificed after ∼2 – 4 hours of imaging. Livers and Lungs were collected and fixed in 10% formalin for 7 days. Samples were embedded in paraffin, cut into 5 µm sections, stained for hematoxylin and eosin (H&E), Histoserv,Inc. The pathology slides were reviewed by Dr. Victoria Hoffmann, Division of Veterinary Resources, Office of the Direction, National Institutes of Health.

### Quantification and statistical analysis

In vivo results represent samples from three to six mice per experimental group. Results represent mean values of at least triplicate samples for ex vivo experiments. Data analysis was performed using Graph-pad Prism 8 (La Jolla, CA, USA) with *t* test or ANOVA. Results are presented as mean ± standard error of mean (S.E.M.). Statistical significance was assessed by p < 0.05 to be considered statistically significant. *p-*values values for each comparison are indicated in the figure legends.

## Supplemental information

Supplemental Information can be found online and includes:

- Figure S1 (related to Fig. 1)
- Figure S2 (related to Fig. 3)
- Figure S3 (related to Fig. 6)
- Figure S4 (related to Fig. 6)
- Figure S5 (related to Fig. 6)
- Figure S6 (related to Fig. 7)
- Table S1 (related to Fig. 3)
- Table S2 (related to Fig. 3)
- Video 1 (related to Fig. 2)
- Video 2 (related to Fig. 2)
- Video 3 (related to Fig. 6)
- Video 4 (related to Fig. 6)
- Video 5 (related to Fig. 6)
- Video 6 (related to Fig. S5)
- Video 7 (related to Fig. 7)
- Video 8 (related to Fig. 7)
- Video 9 (related to Fig. 7)

## Acknowledgements

The authors thank Owen Schwartz of the NIAID Biological Imaging Facility for technical assistance with the microscopy, Larry Lantz of the NIAID Custom Antibody Facility for assistance in conjugating Alexa Fluor fluorochromes to antibodies, Kathleen Harrison for maintaining the G184S mice, Victoria Hoffmann for reviewing the pathology slides, and Dr. Anthony Fauci for long standing encouragement. This work was supported by Intramural Research Program of National Institute of Allergy and Infectious Diseases.

## Author Contribution

SLY, OK, JK, and CP performed the imaging experiments. IYH performed the flow cytometry and migration experiments; and managed the mice. SLY, IYH and JHK designed the experiments. SLY and JHK wrote the manuscript and JHK oversaw the project.

