## Supplement Figures and Tables for "Unrestrained Gα_i2_ Signaling Disrupts Normal Neutrophil Trafficking, Aging, and Clearance"

### Regulator of G Protein Signaling Protein/Gαi2 Interactions Enable Normal Neutrophil Trafficking, Aging, and Clearance

**SUPPLEMENTS**

Figures S1-S6.
Tables S1 & S2
Videos 1-9

**Figure S1 (related to Fig. 1).**

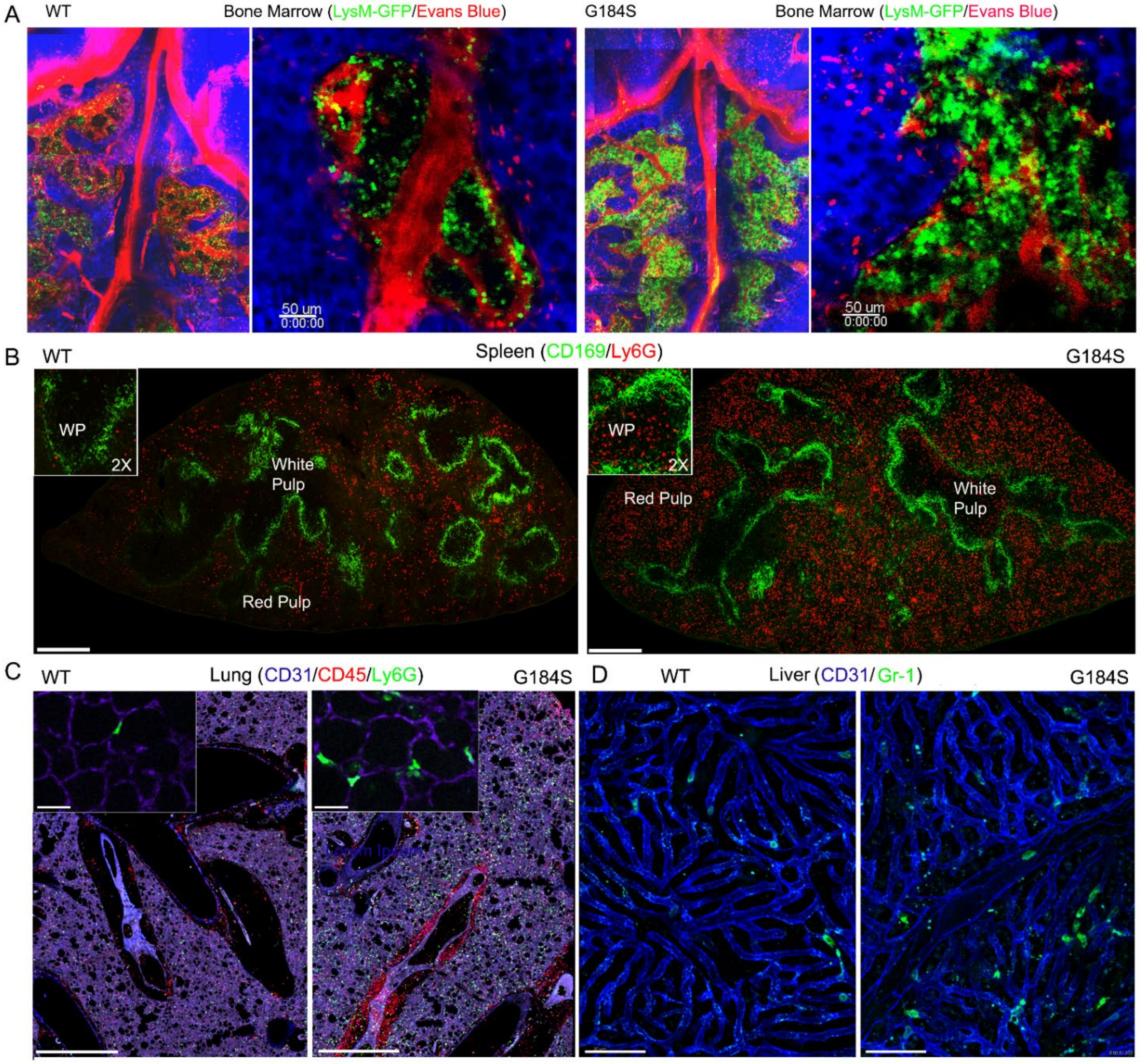


**Figure S1. Identifying WT and G184S Neutrophil Location in the BM, Spleen, Lung, and Liver via Microscopy.** (**A**) Two-photon (2P) microscopy images of skull BM in WT vs G184S mice with LysM-GFP neutrophils (green) and Evans Blue labeled blood vessels (red). Right panels are stitched 2P images spanning the BM niches. Right panels show BM niches in higher magnification during 2P intravital imaging experiments. Results are representative from using WT (n = 5) and G184S (n = 5) mice. Scale bar = 50 µm. (**B**) Representative confocal microscopy images of WT (left panel) vs G184S (right panel) BM reconstituted mice spleens. CD169 immunostaining (green) separates areas of white pulp and red pulp with Ly6G stained neutrophils (red). The larger stitched confocal images show the overall spleen. Scale bar = 400 µm. The insert images show a higher magnification (2X) focusing onto the white pulp area. (**C**) Representative confocal live lung section images of WT (left panel) vs G184S (right panel) BM reconstituted mice. Pulmonary vasculatures (purple) are identified with CD31 immunostaining, CD45 and Ly6G immunostained for leukocytes (red) and neutrophils (green), respectively. The larger stitched confocal images show the overall lung section. Scale bar = 1000 µm. The insert images show a higher magnification focusing into the alveoli. Scale bar = 30 µm. (**D**) Representative confocal intravital microscopy (IVM) liver images of WT (left panel) vs G184S (right panel) BM reconstituted mice. The liver sinusoids (blue) are outlined by CD31 immunostaining, while the neutrophils (green) are identified by Gr-1 staining. Scale bar = 60 µm. (**B**, **C**, **D**) Results are representative images from WT (n = 3) and G184S (n = 3) BM reconstituted mice for each study.

**Figure S2 (related to Fig. 3).**

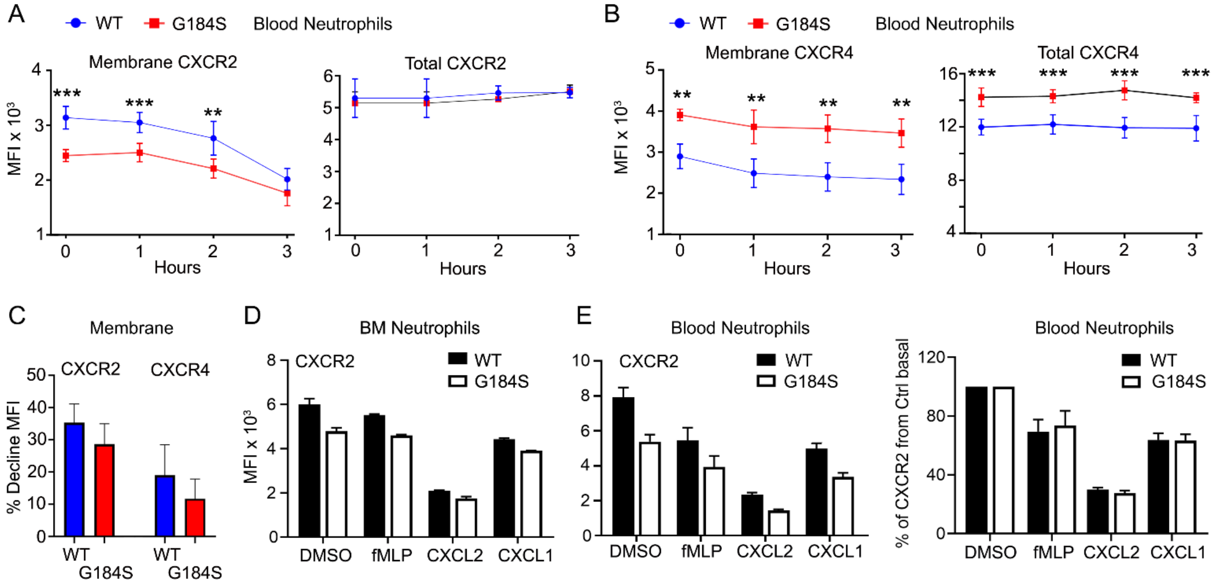


**Figure S2. CXCR4, CXCR2, and CD62L Expression Levels of WT vs G184S Neutrophils.** (**A**) Flow cytometry results measuring membrane (left) and total (right) CXCR2 expression levels of WT vs G184S blood neutrophils in KRH3955 treated mice over 3 hours. (**B**) Results of membrane (left) and total (right) CXCR4 expression levels of WT vs G184S blood neutrophils in KRH3955 treated mice over 3 hours measured by flow cytometry. (**C**) Flow cytometry data showing the final % decline from basal control of membrane CXCR2 and CXCR4 of WT vs G184S blood neutrophils by 3 hours post KRH3955 treatment. (**D, E**) Flow cytometry results measuring membrane CXCR2 expression levels of WT vs G184S neutrophils from BM (**D**) and blood (**E**) exposed to fMLP, CXCL2, and CXCL1 for 30 minutes. Final % decline from basal control of membrane CXCR2 of WT vs G184S blood neutrophils post fMLP, CXCL2, and CXCL1 treatment measured by flow cytometry. All results are from n = 3 separate experiments done in triplicates using n $\geq$ 3 1:1 (WT:G184S) BM reconstituted mice for each study. Statistics: data are means $\pm$ SEMs, then analyzed using unpaired Student's *t*-test comparing G184S with WT. $^{**}p < 0.005$ and $^{***}p < 0.0005$.

**Figure S3 (related to Fig. 6).**

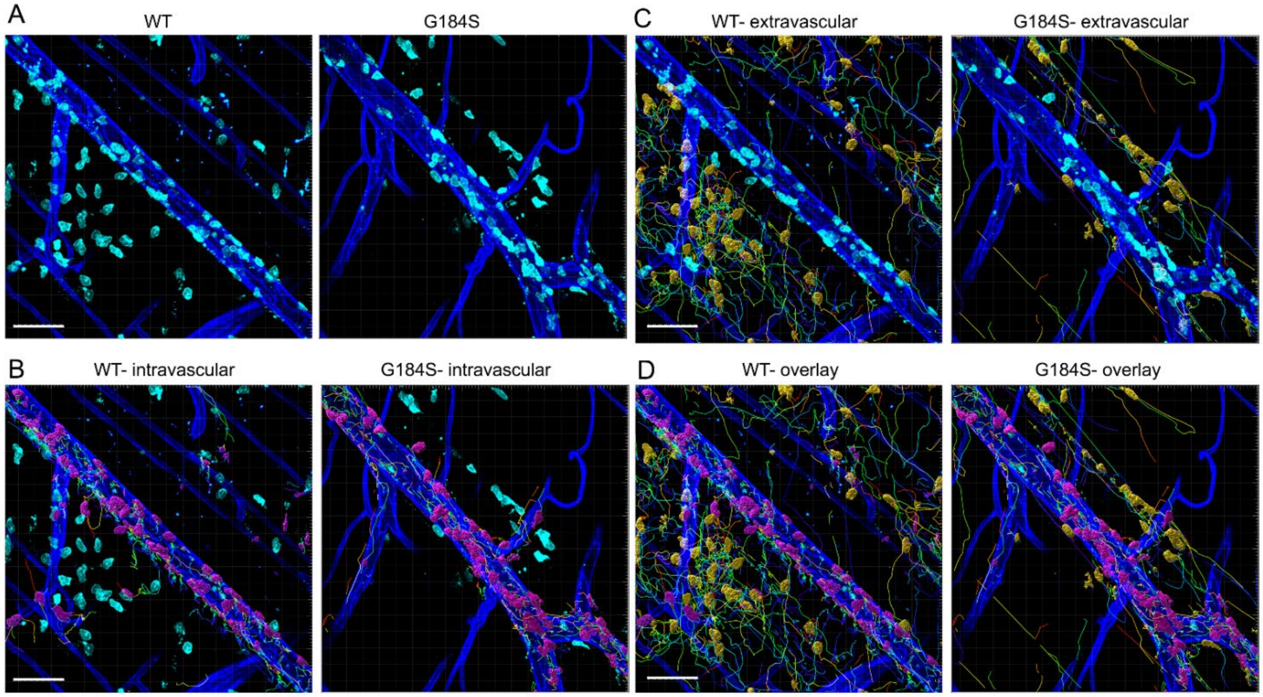


**Figure S3. Distinguishing between Intravascular and Transmigrated Neutrophils.** (**A**) Representative confocal IVM images showing *in vivo* WT (left) vs G184S (right) neutrophil migration in cremaster muscle after 90 minutes of IL-1β (i.v.) and AMD3100 (i.p.) treatment. Neutrophils (cyan) and blood vessels (blue) are visualized by immunostaining with fluorescently labeled Gr-1 and CD31 antibodies, respectively. An example of dividing neutrophils from collected confocal IVM data (**A**) into separate populations of (**B**) intravascular (attached) or (**C**) transmigrated extravascular (free) neutrophils via Imaris software analysis. Using Imaris surface tool, the intravascular neutrophils are set to associate with CD31 staining (attached) and created a new surface (magenta) as identifier (**B**). Transmigrated neutrophils are set to have no association with CD31 staining (free) and created a new surface (yellow) as identifier (**C**). (**D**) After the 2 surfaces are created, neutrophils can be tracked and divided into separate groups (attached vs free) for data analysis. Scale bar = 60 μm.

**Figure S4 (related to Fig. 6).**

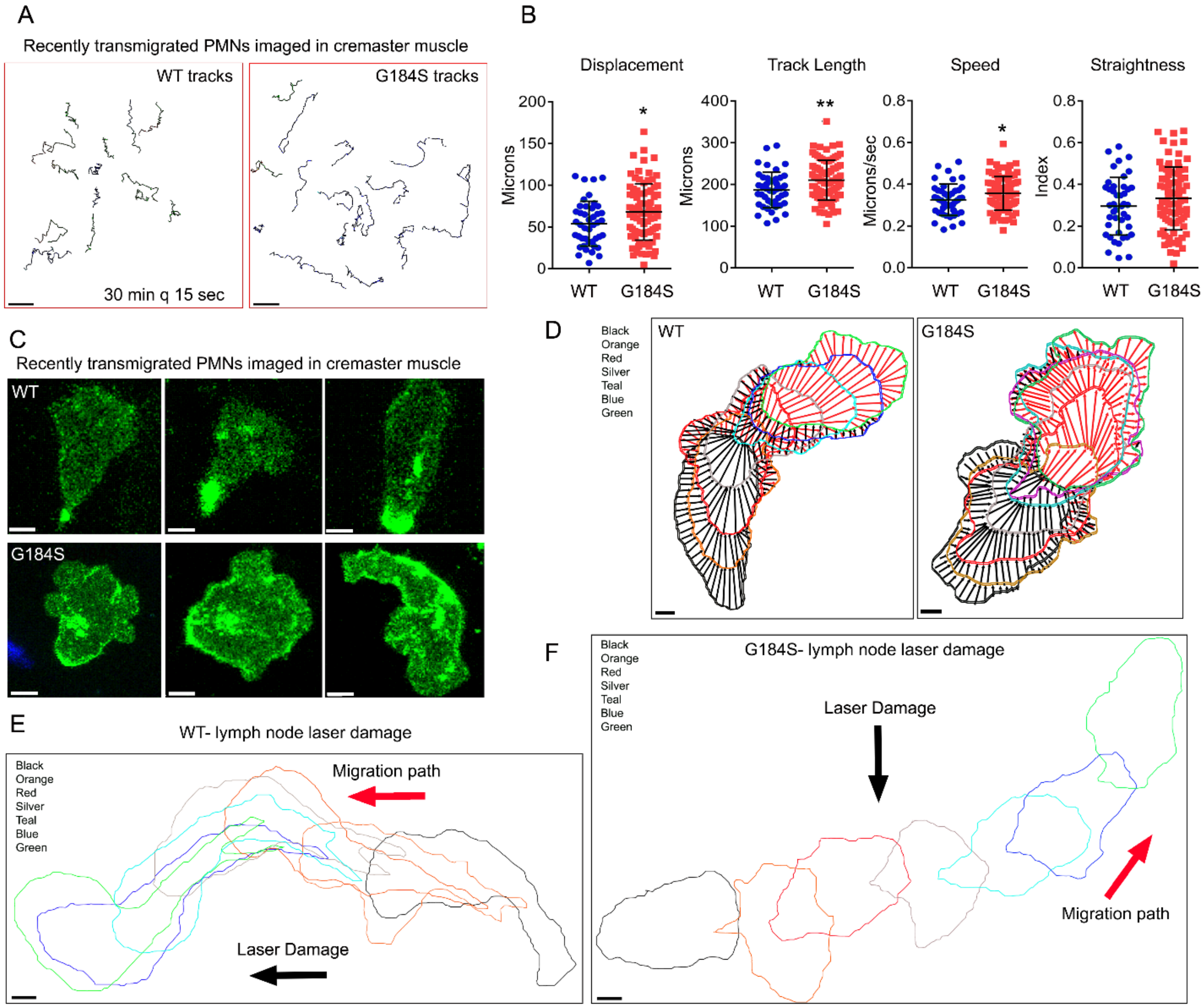


**Figure S4. *In Vivo* Interstitial Migration of WT versus G184S Neutrophils.** (**A**) Representative tracks showing *in vivo* WT (left) vs G184S (right) neutrophil interstitial migration of WT and G184S BM reconstituted mice in the cremaster muscle after IL-1β (i.v.) and AMD3100 (i.p.) treatment. Neutrophils from 4D confocal IVM experiments are tracked for ~30 minutes via Imaris. Scale bar = 30 μm. (**B**) Quantification results for displacement, track length, speed, and straightness from tracking interstitial WT vs G184S neutrophils over 30 minutes from Imaris analysis. (**C**) Representative confocal IVM images shows morphology of individual Gr-1 immunostained WT (top panels) vs G184S (bottom panels) neutrophil migrating in the cremaster muscle interstitium. Scale bar = 3 μm. (**D**) Representative images from Imaris analysis show *in vivo* leading and trailing edge organization of a single WT (left) vs single G184S (right) neutrophil during interstitial migration. Position changes of the neutrophils (1 frame per 25 second) are represented via different colors, with "Black" as the start. Scale bar = 3 μm. (**E, F**) Representative Imaris analysis images of a single WT (**E**) vs single G184S (**F**) neutrophil interstitial migration in mice inguinal lymph node (LN) after laser damage. The black arrows indicate direction of site of laser damage, and the red arrows show the direction of neutrophil migration path. Neutrophil position changes (1 frame per 25 second) are represented via different colors, with "Black" as the start. Scale bar = 3 μm. (**A**, **B**, **C**, **D**) Results are from WT (n = 5) and G184S (n = 5) BM reconstituted mice with 90 min treatment of IL-1β and AMD3100. (**E**) Results are from WT (n = 3) and G184S (n = 3) BM reconstituted mice after 2P laser damage within the LN. Statistics: data are means ± SEMs, then analyzed using unpaired Student's *t*-test comparing G184S with WT. * p < 0.05 and **p < 0.005.

**Figure S5 (related to Fig. 6).**

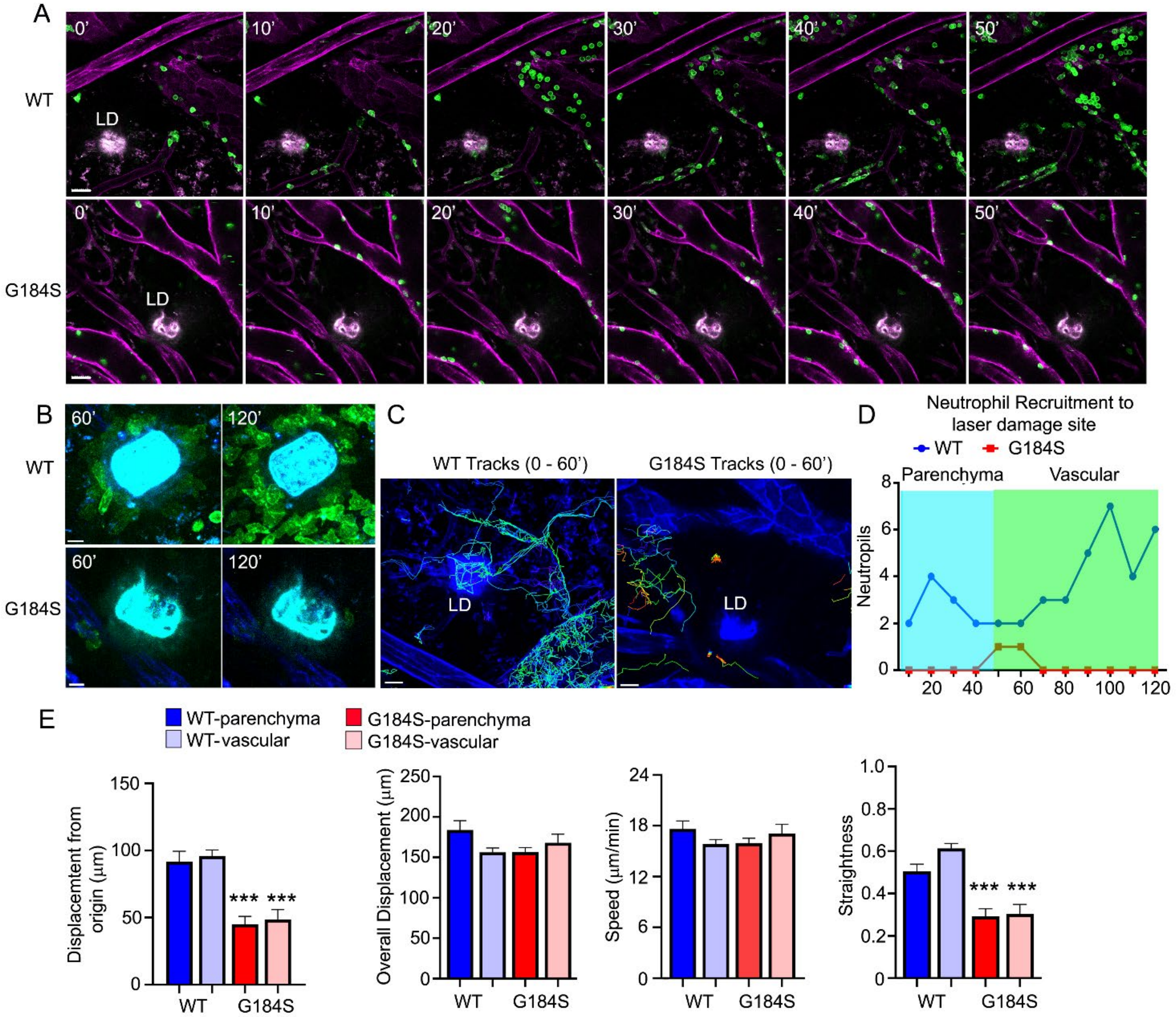


**Figure S5. *In Vivo* Recruitment of WT vs G184S Neutrophils during Sterile Injury.** (**A**) Representative confocal IVM images of *in vivo* WT (top panels) vs G184S (bottom panels) neutrophil directed migration of WT and G184S BM reconstituted mice in inguinal lymph node (LN) after 2P laser damage. Images of Gr-1 immunostained neutrophils (green) migrating within the LN from 0 – 50 minutes after laser damage (LD) are shown. LN vasculatures (magenta) are labeled by CD31 immunostaining. Scale bar = 30 µm. (**B**) Representative confocal IVM images of WT (top panels) vs G184S (bottom panels) neutrophils migrating around LD site (cyan) from **A** at a higher magnification at 60 and 120 minutes after LD damage. Scale bar = 10 µm. (**C**) Representative confocal IVM images showing tracking of WT (left) and G184S (right)

neutrophils around the LD site from **A** for 60 minutes after LD. The multiple tracks are generated by IMARIS imaging software. Scale bar = 20 μm. (**D**) Quantification of accumulating WT vs G184S neutrophils around the LD site from 0 – 120 minutes after LD damage. Migrating cells are divided into either parenchyma or vascular neutrophils. Results are representative from imaging 1 WT and 1 G184S bone marrow reconstituted mouse. (**E**) Displacement from origin, overall displacement, speed, and track straightness for WT (parenchyma, vascular) vs G184S (parenchyma, vascular) neutrophils are shown. Results are from WT (n = 3) and G184S (n = 3) BM reconstituted mice all tracked over 60 minutes after LD. Statistics: data are means $\pm$ SEMs, then analyzed using unpaired Student's *t*-test comparing parenchyma-G184S with parenchyma-WT and vascular-G184S with vascular-WT. ***p < 0.0005.

**Figure S6 (related to Fig. 7).**

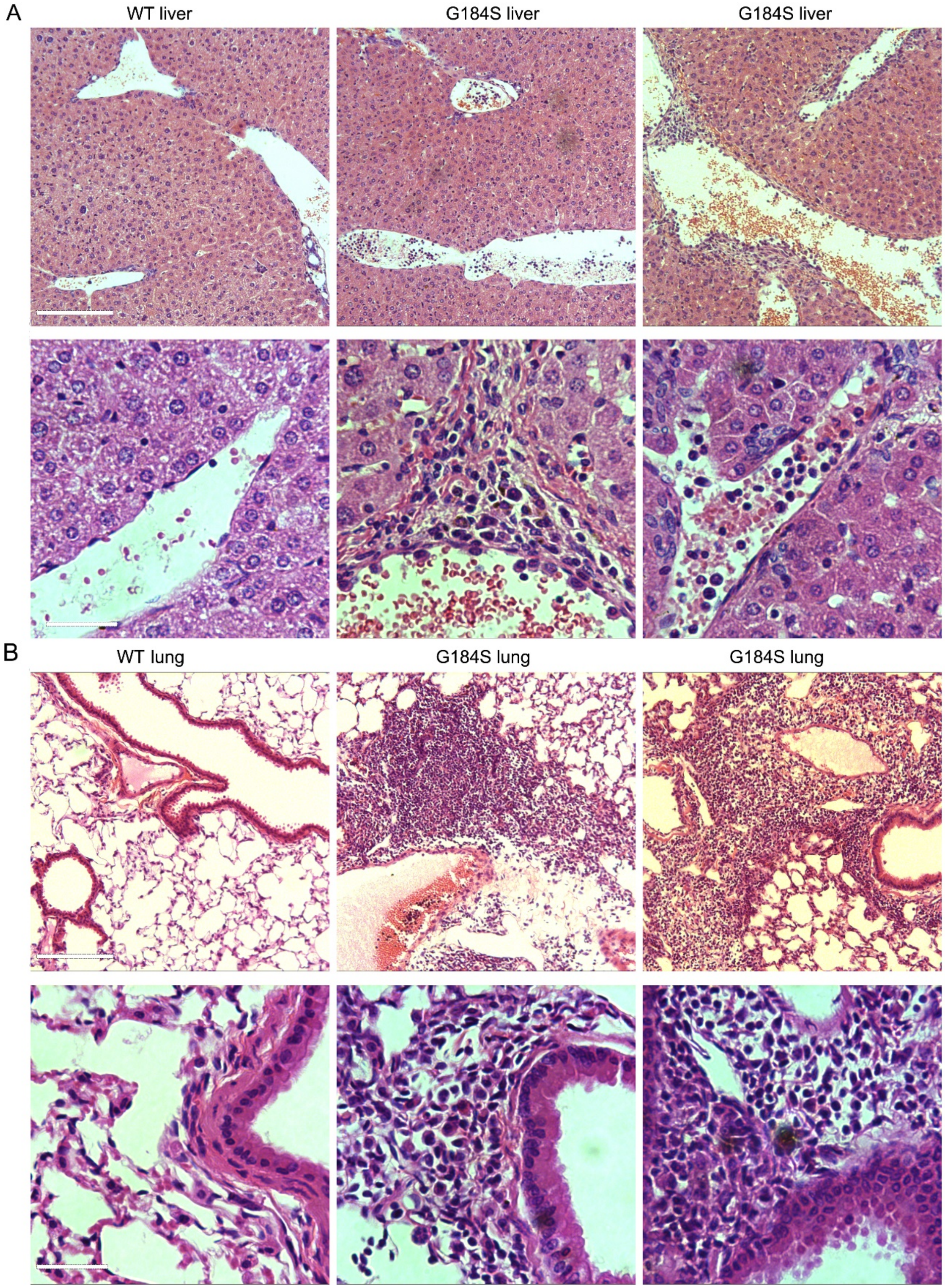


•

**Figure S6. Hepatic and Pulmonary Neutrophil Infiltration in WT vs G184S after ConA Injection.** (A) Representative liver images from histopathological examination comparing 1 WT to 2 G184S BM reconstituted mice after 3 hrs. ConA injection. 10X (top panels) vs 40X (bottom panels) magnification. 10X, scale bar = 150 µm; 40X, scale bar = 30 µm. (**B**) Representative lung images after histopathological examination comparing 1 WT to 2 G184S BM reconstituted mice after 3 hrs. ConA injection. 10X (top panels) vs 40X (bottom panels) magnification. 10X, scale bar = 150 µm; 40X, scale bar = 30 µm. Hematoxylin - eosin staining results are from WT (n = 2) and G184S (n = 3) BM reconstituted mice.

**Table S1 (related to Figure 3).**

Table S1

| | | BM Immature | | BM Mature | |
|---|---|---|---|---|---|
| | | WT (CD45.1, n=3) | G184S (CD45.2, n=3) | WT (CD45.1, n=3) | G184S (CD45.2, n=3) |
| CXCR4 | extracellular | 160.0 ± 0.58 | 211.0 ± 4.04 | 108.5 ± 1.44 | 191.5 ± 0.29 |
| | Intracellular | 910.5 ± 59.18 | 937.0 ± 22.52 | 961.5 ± 77.65 | 1130.5 ± 34.35 |
| | Ratio ($MFI_{extra}/MFI_{Intra}$) | 0.18 ± 0.012 | 0.23 ± 0.010 | 0.11 ± 0.011 | 0.17 ± 0.005 |
| CXCR2 | extracellular | 773.5 ± 23.96 | *** 524.5 ± 80.54 | 1940.0 ± 32.33 | *** 1641.5 ± 116.91 |
| | Intracellular | 358.0 ± 25.40 | 421.5 ± 75.34 | 360.5 ± 54.56 | 826.0 ± 101.04 |
| | Ratio ($MFI_{extra}/MFI_{Intra}$) | 2.19 ± 0.224 | *** 1.41 ± 0.465 | 5.68 ± 0.982 | *** 2.09 ± 0.406 |
| CD62L | extracellular | 2118.5 ± 4.91 | *** 1523.5 ± 11.84 | 2904.5 ± 7.22 | 2747.5 ± 29.73 |

**Table S1. Neutrophil CXCR4, CXCR2, and CD62L Expression Levels on Bone Marrow Neutrophils.** Flow cytometry data shows CXCR4, CXCR2, and CD62L expression levels of WT vs G184S BM neutrophils. Neutrophils isolated from bone marrows of 1:1 chimeric mice are further divided into immature and mature subsets. Data are means ± SEMs. Student's *t*-test comparing G184S with WT neutrophils. ***$p < 0.0005$.

**Table S2 (related to Figure 3).**

Table S2

| | | Spleen | | Lung | | Liver | |
|---|---|---|---|---|---|---|---|
| | | WT (CD45.1, n=3) | G184S (CD45.2, n=3) | WT (CD45.1, n=3) | G184S (CD45.2, n=3) | WT (CD45.1, n=3) | G184S (CD45.2, n=3) |
| CXCR4 | extracellular | 163.5 ± 3.18 | 209.7 ± 4.91 | 181.5 ± 2.02 | 150.0 ± 5.20 | 184.0 ± 13.86 | 219.0 ± 16.17 |
| | Intracellular | 1670.0 ± 12.12 | 1623.0 ± 32.91 | 2157.0 ± 55.43 | 2230.5 ± 77.65 | 1242.5 ± 21.65 | 1249.0 ± 113.61 |
| | Ratio ($MFI_{extra}/MFI_{Intra}$) | 0.10 ± 0.003 | 0.13 ± 0.005 | 0.08 ± 0.001 | 0.07 ± 0.005 | 0.15 ± 0.014 | 0.18 ± 0.030 |
| CXCR2 | extracellular | 2014.5 ± 21.07 | *** 925.5 ± 10.10 | 1744.5 ± 43.59 | *** 699.5 ± 14.15 | 797.0 ± 6.35 | *** 310.5 ± 2.60 |
| | Intracellular | 2805.0 ± 20.79 | 2639.0 ± 36.37 | 1638.5 ± 63.80 | 1755.0 ± 12.70 | 1687.0 ± 15.59 | 1379.5 ± 7.79 |
| | Ratio ($MFI_{extra}/MFI_{Intra}$) | 0.72 ± 0.002 | *** 0.35 ± 0.001 | 1.07 ± 0.068 | *** 0.40 ± 0.011 | 0.47 ± 0.008 | *** 0.23 ± 0.003 |
| CD62L | extracellular | 842.0 ± 1.16 | *** 397.5 ± 7.22 | 508.0 ± 22.52 | *** 338.5 ± 1.44 | 591.0 ± 2.31 | *** 261.5 ± 2.60 |

**Table S2. Neutrophil CXCR4, CXCR2, and CD62L Expression Levels on Spleen, Lung, and Liver Neutrophils.** Flow cytometry data shows CXCR4, CXCR2, and CD62L expression levels of WT vs G184S BM reconstituted mice. Neutrophils were isolated from spleen, lung, and liver from 1:1 chimeric mice. Data are means ± SEMs. Student's *t*-test comparing G184S with WT neutrophils. ***$p < 0.0005$.

#### Video Legends

**Video S1 (Related to Figure 2). *In Vivo* Visualization of WT vs G184S Basal Neutrophil Mobility within Murine Bone Marrow Niches.** The multi-photon (MP) intravital microscopy (IVM) movie on the left captures the mobility of WT neutrophils (green) within the skull bone marrow (BM) of a PBS-treated LysM-GFP x WT BM reconstituted mouse. The sequence shows a ~60-min period. The MP-IVM movie on the right shows the mobility of G184S neutrophils (green) within the skull BM of a PBS-treated LysM-GFP x G184S BM reconstituted mouse. The sequence shows a ~55-min period. Blood vessels are outlined by i.v. injection of Evans Blue (red). Images were captured at ~1 frame per 12 s with ~20X magnification.

**Video S2 (Related to Figure 2). Visualizing WT vs G184S Neutrophil Mobility within Stimulated Murine Bone Marrow Niches *In Vivo*.** The MP-IVM movie on the left captures the mobility of WT neutrophils (green) within the skull BM of a CXCL1-treated LysM-GFP x WT BM reconstituted mouse. The sequence shows a ~60-min period. The MP-IVM movie on the right shows the mobility of G184S neutrophils (green) within the skull BM of a CXCL1-treated LysM-GFP x G184S BM reconstituted mouse. The sequence shows a ~43-min period. Blood vessels are outlined by i.v. injection of Evans Blue (red). Images were captured at ~1 frame per 12 s with ~20X magnification.

**Video S3 (Related to Figure 6). *In Vivo* Visualization of WT vs G184S Neutrophil Trafficking in Inflamed Murine Cremaster Muscle Venules.** The confocal IVM movies compared WT (left) and G184S (right) neutrophil trafficking in IL-1β (i.s.) + AMD3100 (i.p.)-treated cremaster muscles with Alexa Fluor 488-anti-Gr1 mAb neutrophils (cyan) and Alexa Fluor 647-anti-CD31 mAb venules (blue). On the left, the movie captures WT neutrophils adhering, crawling, transmigrating through venular walls, and migrating within the interstitial tissue in a WT BM reconstituted mouse. The sequence shows a ~60-min period. The movie on the right shows G184S neutrophils trafficking within and outside the inflamed venules in a G184S BM reconstituted mouse. The sequence shows a ~60-min period. Images were captured at ~1 frame per 30 s with ~25X magnification.

**Video S4 (Related to Figure 6). *In Vivo* WT Neutrophil Transmigration in Inflamed Murine Cremaster Muscle Venules.** At a higher magnification, these confocal IVM movies capture Alexa Fluor 488-anti-Gr1 mAb immunostained WT neutrophils (cyan) undergoing transmigration (TEM) in WT BM reconstituted mice treated with IL-1β (i.s.) and AMD3100 (i.p.). Endothelial cells (ECs) of the venule segments (blue) were immunostained *in vivo* with Alexa Fluor 647-anti-CD31 mAb. The left movie shows a WT neutrophil undergoing TEM. The movie taken from the luminal side shows an incoming WT neutrophil adhering to venule at EC junctions, and squeezing through the EC junction via extensive changes in neutrophil cell shape. The yellow track shows the course of WT neutrophil TEM with the appearance and movement of the yellow sphere indicating the start to end of this process. The sequence shows a ~18-min period. The right confocal IVM movie taken from the luminal side captures 2 WT neutrophils undergoing consecutive TEM (yellow and red tracks) through an opened endothelial pore "hot spot" located on a venule segment. The spheres (yellow and red spheres) appearing on the tracks indicate the start to end of each process. The sequence shows a ~17-min period. Images were captured at ~1 frame per 30 s with ~75X magnification.

**Video S5 (Related to Figure 6). Visualizing Intravascular G184S Neutrophil Behaviors and Transmigration *In Vivo*.** At a higher magnification, these confocal IVM movies capture the intravascular behaviors and TEM of Alexa Fluor 488-anti-Gr1 mAb immunostained G184S neutrophils (cyan) in G184S

BM reconstituted mice treated with IL-1β (i.s.) and AMD3100 (i.p.). The venule segment (blue) are immunostained *in vivo* with Alexa Fluor 647-anti-CD31 mAb. The top left movie shows a G184S neutrophil adhering to the venule segment. The adherent G184S neutrophil eventually dislodged and disappeared into the circulation. The sequence shows a ~18-min period. The top right movie captures an incoming circulating G184S neutrophil adhering to and crawling along the intravascular venular wall, eventually detaching and returns to the circulation. The sequence shows a ~17-min period. The movie on the bottom left shows an incoming circulating G184S neutrophil adhering to the venule, squeezing through the EC wall via substantial changes in neutrophil cell shape. The sequence shows a ~23-min period. The movie on the bottom right captures an incoming circulating G184S neutrophil adhering to and crawling along the intravascular venular wall, eventually undergo TEM. The sequence shows a ~30-min period. The yellow spheres appearing along the tracks indicate the starting and ending of the described processes. Images were all taken from the luminal side and captured at ~1 frame per 30 s with higher magnification (~60 – 75X).

**Video S6 (Related to Figure S5). *In Vivo* Visualization of WT vs G184S Neutrophil Trafficking in Laser Damaged Murine Inguinal Lymph Nodes.** The confocal IVM movies compared WT (left) and G184S (right) neutrophil trafficking within inguinal lymph nodes (LNs) after laser-induced damage (white spot). The LN vasculature (magenta) was visualized with Alexa Fluor 555-anti-CD31 mAb immunostaining, and the Alexa Fluor 488-anti-Gr1 mAb immunostained the WT neutrophils (green) *in vivo*. The left movie captures the result of WT neutrophil trafficking after laser-induced damage within the LN of a WT BM reconstituted mouse. The WT neutrophils accumulate within high endothelial venules (HEVs) near the laser damage site and eventually transmigrate into the LN tissue and travel towards the damage site. The sequence shows a ~60-min period. On the right, the movie shows the G184S neutrophils trafficking within an inguinal LN of a G184S BM reconstituted mouse after laser damage. The sequence shows a ~60-min period. Laser damage was induced inside the LN parenchyma by applying 50% of the MP laser power at zoom 25 for 3 s. Images were taken immediately after laser damage at ~1 frame per 30 s with ~ 25X magnification.

**Video S7 (Related to Figure 7). Visualizing WT vs G184S Neutrophil Trafficking in Inflamed Murine Liver *In Vivo.*** The confocal IVM movie on the left shows the result of WT neutrophil trafficking and activity in the liver of a WT BM reconstituted mouse with ConA-induced inflammation. The WT neutrophils (green) and liver sinusoidal vasculature (blue) are immunostained *in vivo* with Alexa Fluor 488-anti-Gr1 mAb and Alexa Fluor 647-anti-CD31 mAb, respectively. The sequence shows a ~40-min period. On the right, the movie captures the result of G184S neutrophil trafficking and activity in the liver of a G184S BM reconstituted mouse with ConA-induced inflammation. The sequence shows a ~40-min period. Images were captured at ~1 frame per 30 s with ~25X magnification.

**Video S8 (Related to Figure 7). *In Vivo* Visualization of WT vs G184S Neutrophil and Platelet Interactions in Inflamed Murine Liver.** On the left, the confocal IVM movie shows the result of WT neutrophil (green) and WT platelets (red) immunostained with Alexa Fluor 488-anti-Gr1 mAb and DyLight 649-anti-GP1bβ mAb, respectively, during liver inflammation. The neutrophils and platelets traffic and interact within the Alexa Fluor 555-anti-CD31 mAb immunostained liver sinusoid (violet) of a WT BM reconstituted mouse with ConA-induced inflammation. The adherent neutrophils interact with circulating platelets and platelets aggregates to form clots. The sequence shows a ~30-min period. The right confocal IVM movie captures the result of G184S neutrophil and platelets interacting within the liver sinusoid of a G184S BM reconstituted mouse with ConA-induced inflammation. The platelets interact with adherent neutrophils, formed platelet aggregates, and larger platelet clots within the sinusoid. The sequence shows a ~30-min period. Images were captured at ~1 frame per 30 s with ~50X magnification.

**Video S9 (Related to Figure 7).** ***In Vivo*** **Neutrophil Fragmentation and Netosis in Inflamed Murine Liver.** The left confocal IVM movie shows the fragmentation of an Alexa Fluor 488-anti-Gr1 mAb immunostained G184S neutrophil (green) within a liver of a G184S BM reconstituted mouse with ConA-induced inflammation. The sinusoid vasculature (blue) are immunostained with Alexa Fluor 647-anti-CD31 mAb. In the middle, the neutrophil crawled along the sinusoid, elongates, and breaks off a small fragment. The sequence shows a ~26-min period. On the right, the confocal IVM movie captures a G184S neutrophil undergoing netosis within the liver sinusoid during ConA-induced inflammation. The sequence shows a ~40-min period. Images were captured at ~1 frame per 30 s with ~65X magnification.
